# A low-dimensional, generalizable encoding manifold for auditory cortex

**DOI:** 10.64898/2026.08.30.747962

**Authors:** Satyabrata Parida, Jereme C. Wingert, Jonah D. Stickney, Stephen V. David

## Abstract

Neural populations in auditory cortex (AC) perform sensory computations that support stimulus category decoding and flexible behavior. The underlying geometry and generalizability of these computations for natural stimuli remain poorly understood, particularly at the single-neuron level, where neurons display wide-ranging tuning specificity and temporal acuity. To address this gap, we developed ACNet, a foundation model of cortical sound encoding, to predict the time-varying activity of >3000 neurons in AC of ferrets. Training data included 42 hours of natural sounds spanning over 100 categories and were collected from multiple recording sites across multiple animals. The model achieved state-of-the-art response prediction accuracy. Model activity was succinctly captured by a low-dimensional neural manifold, which generalized (>80% of variance) across animals. Analysis of ACNet activations revealed the emergence of rate-based, sparse sound coding across layers, a prominent feature of the auditory cortex. These transformations were concomitant with the emergence of more accurate and neurally aligned auditory category decoding, even though ACNet was not explicitly trained to categorize sounds. The model also revealed tuning differences between anatomically distinct cell types. Taken together, our results demonstrate that foundation models of sensory systems can reveal generalizable computations by large neural populations.

## Introduction

Accurate computational models are critical for assessing our understanding of neural sensory systems. Computational models that account for neural functions enable *in silico* hypothesis testing, refinement of experimental design, and applications that align with neural principles (Kriegeskorte, 2015; Kriegeskorte & Douglas, 2018; Richards et al., 2019; Sadagopan et al., 2023; Saxe et al., 2021). Accurate encoding models, particularly at the level of the cortex, must capture functional properties across diverse neural populations and across a vast range of sensory contexts (Carandini et al., 2005; David, 2018; E. Y. Wang et al., 2025; Wu et al., 2006). Traditional encoding models rely on a phenomenological approach, either hand-engineered to match limited data, e.g., Gabor filter models for spectrotemporal or spatiotemporal modulation (Adelson & Bergen, 1985; Chi et al., 2005; Fishbach et al., 2003; Harper et al., 2016; Jones & Palmer, 1987), or derived theoretically (Marĉelja, 1980; Młynarski & McDermott, 2018). An alternative to hand-engineered approaches are task-optimized models, based on artificial neural networks trained on tasks such as object classification (Bakhtiari et al., 2021; Francl & McDermott, 2022; Kell et al., 2018; Saddler et al., 2021; Yamins et al., 2014). These task-optimized models have been able to account for novel, nonlinear coding properties (Cichy et al., 2016; Kell et al., 2018; Khatami & Escabí, 2020; Yamins et al., 2014), but they use computations that diverge from those used by biological systems (Feather et al., 2023; Muzellec & Kar, 2026; Tuckute et al., 2023; Weerts et al., 2022). This divergence is particularly notable in the regime of spike timing, which often carries precise information in the brain, notably within the auditory cortex (Joris et al., 2004; Lu et al., 2001; Pennington & David, 2023), but is not typically assessed for task-optimized models designed to account for time-averaged neural activity.

Recent advances in large-scale data collection, combined with deep learning, have paved the way for a new class of neural network model trained directly on neural activity (Cadena et al., 2019; d’Ascoli et al., 2026; Richards et al., 2019; E. Y. Wang et al., 2025). These foundation models are often fit to large, diverse datasets, including data from different subjects (Drakopoulos et al., 2025; Dupré la Tour et al., 2025; E. Y. Wang et al., 2025). This approach provides access to a *sensory encoding manifold*, the geometry of neural population activity evoked by wide-ranging sensory inputs (Kriegeskorte & Wei, 2021; Sabesan et al., 2023). A primary goal of this approach is to capture core encoding properties that generalize across subjects, species, and sensory contexts (P.-H. (Cameron) Chen et al., 2015; Güçlü & van Gerven, 2017; Haxby et al., 2011; Sabesan et al., 2023; Schneider et al., 2023; Yang et al., 2026). However, it remains unclear whether the encoding manifold framework can reveal canonical computations that generalize across subjects and sensory contexts while also capturing the heterogeneity of single-neuron selectivity, particularly at the temporal resolution needed to resolve the auditory system’s fast dynamics.

To bridge this gap between generalizable computations and the substantial heterogeneity of single neuron selectivity, we present **ACNet**, a foundation model trained to predict the activity of 3,124 single AC neurons with 10-ms temporal precision. To ensure generalizability, training data included over 42 hours of natural sounds and data from multiple recording sites across multiple subjects. ACNet matches the performance of state-of-the-art CNNs trained on individual recording sites, while establishing a low-dimensional neural encoding manifold (referred to as the manifold hereafter for brevity) that generalizes with a high degree of representational similarity across animals. Manifold activity was concomitant with an emergent representation of auditory categories, even though the encoding model was not explicitly trained to classify sounds. Activations of the manifold also reproduced multiple hallmark features of the ascending auditory pathway (Atencio et al., 2012; Chechik et al., 2006; Lu et al., 2001). Together, these findings support the idea that a large-scale characterization of neural sound encoding can provide a generalizable model of auditory cortical function.

## Results

### ACNet predicts natural sound-evoked activity in auditory cortex

To sample neural activity comprehensively from the auditory cortex (AC), we used high-channel count electrodes to record single-unit activity from multiple recording sites in primary (A1) and secondary (PEG) fields of passively listening, head-fixed ferrets (3124 units, 62 sites, 4 animals; Figure 1). During recordings, we presented sequences of natural sound samples (50-800 ms duration) drawn from wide-ranging categories (Wingert et al., 2026). Individual recordings included 15-150 min of stimuli (mean=41 min), which were sampled differentially across experiments for a total of 42 hours (20.5 hours unique) of natural sounds.

**Figure 1.**
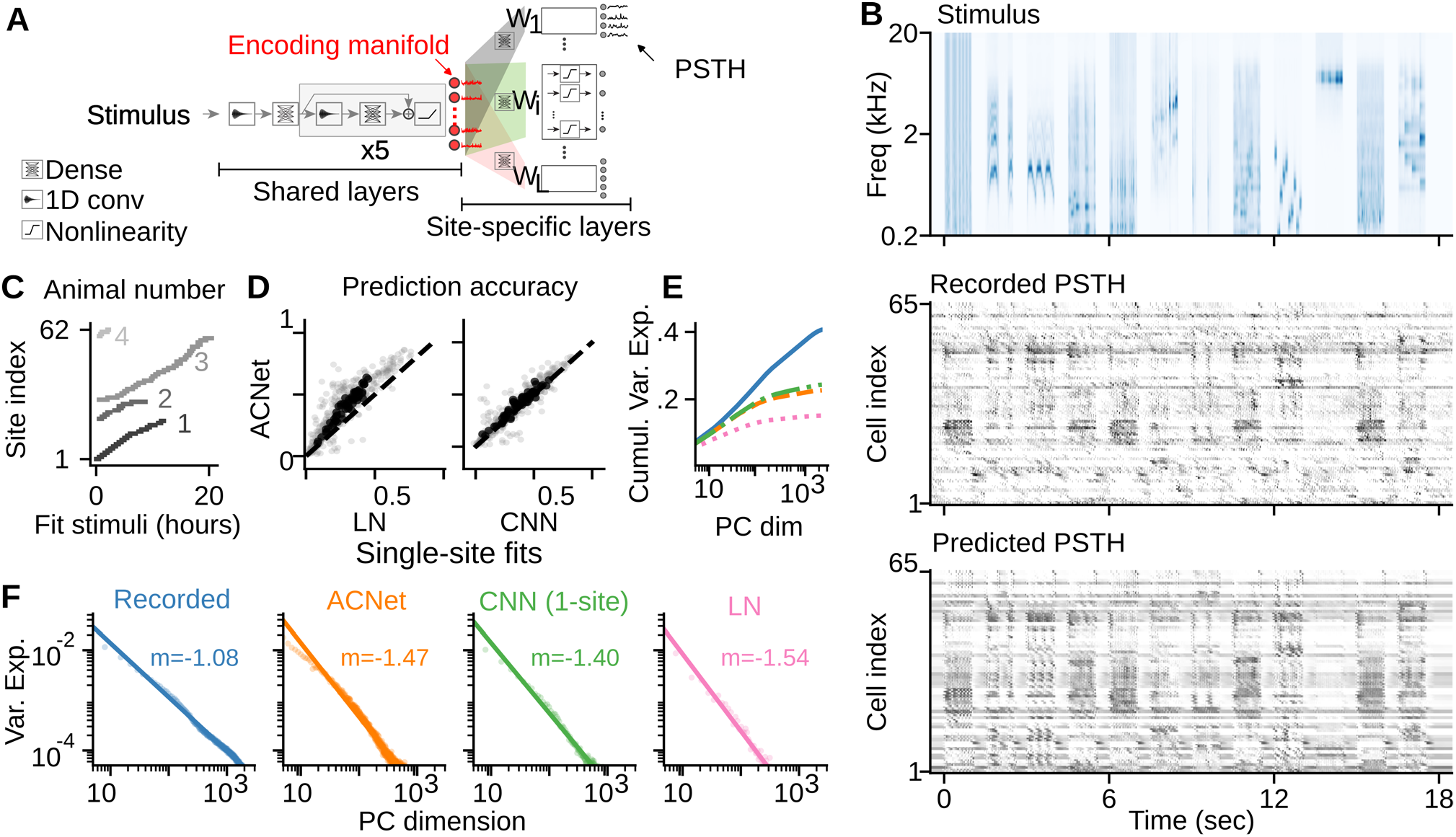
ACNet can accurately predict auditory cortical responses across animals. **(A)** ACNet is a residual CNN with multiple heads, each corresponding to a different recording site. The model was trained simultaneously to predict neural responses for all sites from the spectrogram of natural sound stimuli. **(B)** Example stimulus spectrogram and corresponding actual and predicted neural activity for validation data from one recording site, plotted one row per neuron. **(C)** Sequences of natural sounds were played to passively listening, awake, head-fixed ferrets (3124 cells, 62 recording sites, 4 animals). Stimuli used for model training were generally distinct across sites [41 (mean) ± 24 (sd) min/site, 42 hours total and 20.5 hours unique, cumulatively across sites] and played without repetition. A standard set of stimuli for model validation (108-s duration) was repeated 10 times for each site. **(D)** Model accuracy, measured as the correlation coefficient between actual and predicted PSTH responses for the validation stimuli, was greater for ACNet than for linear-nonlinear (LN) models (median difference=0.09, p<1e-16, one-sided Wilcoxon signed-rank test) and slightly higher than that for state-of-the-art CNNs fit separately for each recording site (median difference=0.02, p<1e-16, one-sided Wilcoxon signed-rank test). Gray dots compare model accuracy for single neurons; black for recording site means. **(E)** Cross-validated cumulative variance explained for increasing numbers of principal components (PCs) computed from time-varying neural activity and from predictions by ACNet, single-site CNN, and LN models. Colors match labels in F. **(F)** Fraction of variance explained for individual PCs of actual and model activity. Solid lines are power-law functions fit to dimensions 5-400 (m: power-law exponent).

To map the relationship between stimulus and neural response, a convolutional neural network (CNN) with an encoder-decoder architecture was trained to predict the responses of neuronal populations to the sound spectrogram (Figure 1A). The encoder was composed of six convolutional blocks with residual connections (He et al., 2016). The decoder was multi-headed, with each head predicting the activity of a single recording site. Decoding heads were memoryless, performing a linear weighting of the encoder output, followed by a static nonlinearity. We refer to this model as ACNet. ACNet outperformed traditional linear-nonlinear (LN) models (Figure 1D) in predicting held-out validation data, replicating LN versus CNN comparisons in the auditory (Pennington & David, 2023) and visual systems (Cadena et al., 2019; E. Y. Wang et al., 2025). ACNet performed similarly to independent CNNs fit separately for each recording site (Figure 1D), suggesting that the encoder captures a complete basis set of transformations for the entire neuronal population.

Studies in visual cortex have shown that neural population responses to natural stimuli are high dimensional and optimally balance the redundancy and efficiency of the neural code (Stringer et al., 2019). To test whether this optimal geometry holds true in AC, we quantified the variance and dimensionality of the held-out validation data using cross-validated Principal Component Analysis (cvPCA). This approach provides unbiased estimates of the variance explained by each dimension by fitting PCs on half of the neural dataset and measuring variance with the other half of the neural dataset (noise ceiling) or with model predictions of the neural data. Comparing model performance (e.g., ACNet vs. single-site CNN) reveals how much of the unique neural variance each model captures and how this variance is distributed across PC dimensions. Consistent with their higher prediction accuracy, ACNet and single-site CNNs explained 55.8% and 59.8% of the explainable population variance, respectively, substantially outperforming the LN model (37.3%) (Figure 1E).

To estimate how variance scales across dimensions in the neural population, a power-law function was fit to the cvPCA eigenspectrum (variance explained curve; Figure 1F). Mathematically, an exponent (slope) close to -1 represents optimal 1/f processing by neural populations balancing efficiency and redundancy, whereas more negative (or less negative) exponents reflect redundant (or efficient) processing. The exponent for our AC data was -1.08, which is remarkably similar to the value of -1.04 measured in mouse visual cortex (Stringer et al., 2019). This result supports the theory that this high-dimensional population geometry is a fundamental property of optimal cortical processing, generalizing across sensory modalities and species. LN models had the steepest slope, followed by ACNet and single-site CNNs. The lower variance explained and steeper dimensionality of ACNet relative to single-site CNNs, paired with the slightly higher prediction accuracy of ACNet, may reflect the trade-off that ACNet better captures the dominant tuning properties that generalize across sites, but partly filters out idiosyncratic neuronal properties within a site.

### ACNet identifies a generalizable encoding manifold of auditory cortex

The output of the ACNet encoder, or the *encoding manifold*, provides a basis set of stimulus transformations that predict the activity across the entire neuronal population. We surmised that if this basis set is comprehensive for AC, then it should generalize across animals (Figure 2A). To test this idea, we used representational similarity analysis (RSA) to perform a nonparametric comparison of the encoding layer output between animals (Kriegeskorte et al., 2008). We trained separate instances of the ACNet model for each animal and then played the validation stimulus to obtain activations for the encoder layer of each animal-specific model. The Euclidean distance between activation for each stimulus was used to compute the representational dissimilarity matrix (RDM; Figure 2B), which quantifies the pairwise dissimilarity of each stimulus’s representation within each animal’s manifold. To evaluate representational similarity across animals, we computed the Pearson’s correlation coefficient between their respective RDMs.

**Figure 2.**
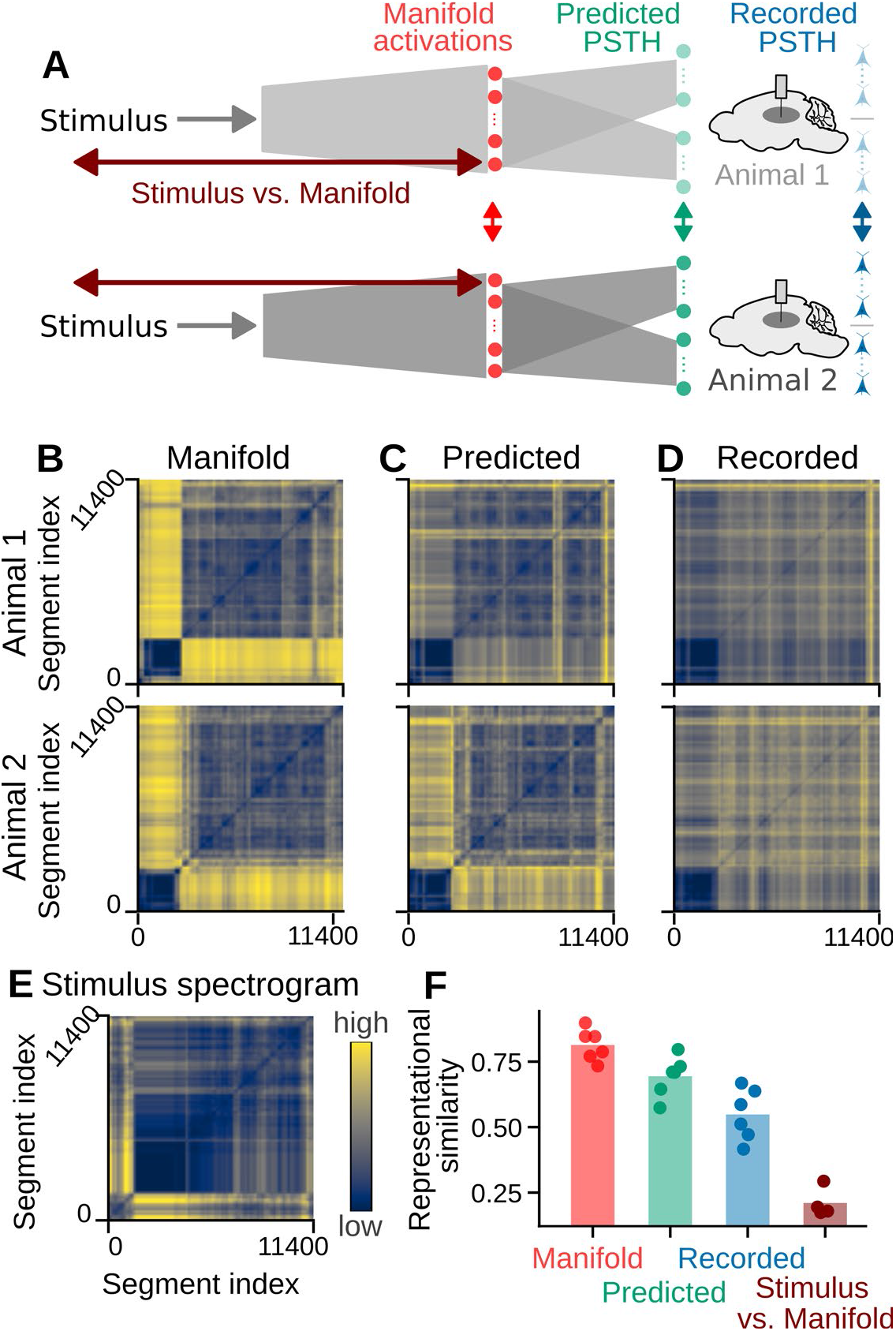
Auditory cortical manifold generalizes across animals. **(A)** Schematic of the three model stages to which Representational Similarity Analysis (RSA) was applied. **(B)** Representational dissimilarity matrices (RDMs) for the manifold (output of the encoding layer evoked by test stimuli) from ACNet trained separately with data from two different animals. Each matrix value indicates the difference between activation at different 10-ms stimulus time bins. Cooler (warmer) colors indicate smaller (larger) Euclidean distance. **(C)** RDM for predicted single-neuron activity for the same animals. **(D)** RDM for recorded neural activity for the same animals. **(E)** RDM for the spectrogram of the test stimulus. **(F)** Representational similarity (i.e., correlation coefficient between RDMs) between animals for manifold activations (0.82±0.06) and predicted (0.69±0.08) and recorded single unit PSTHs (0.56±0.10), as well as between manifold and stimulus spectrogram (0.22±0.05). Manifold similarity across animals was significantly greater than predicted PSTH similarity (difference = 0.13, 95% CI [+0.10, +0.15], measured with 1000 resamples of 1-s stimulus blocks), recorded PSTH similarity (difference = 0.26, 95% CI [+0.21, +0.32]), and manifold-stimulus similarity (difference = 0.60, 95% CI [+0.56, +0.63]). CI: confidence interval.

Representational similarity between manifold layer activations (mean *r*=0.82) was greater than between the predicted (*r*=0.69) and actual (*r*=0.56) single neuron responses to the same stimuli (Figure 2F). This greater similarity is consistent with the manifold being a basis set that supports the variable tuning in individual neurons, which were sampled differently across animals (Figures 2C and 2D). Furthermore, this alignment is not a trivial consequence of low-level stimulus statistics, as the manifolds exhibited low similarity to the raw gammatone spectrogram (*r*=0.22, Figure 2E). The strong representational similarity of animal-specific manifolds indicates that the ACNet manifold does not merely fit individual datasets, but instead uncovers a conserved, canonical set of sensory transformations that generalize across subjects (Figure 2F).

### ACNet manifold embedding supports neurally aligned category decoding

A critical task the auditory system must accomplish is sound categorization, i.e., classifying acoustic objects according to their behavioral meaning. Modern task-optimized deep neural networks (DNNs) trained explicitly to perform sound classification can achieve remarkable accuracy (K. Chen et al., 2022; S. Chen et al., 2022; Gong et al., 2021; Kong et al., 2020). While these models can replicate human perceptual confusions in some listening conditions (e.g., in background noise or spectrotemporal filtering) (Alavilli & McDermott, 2026; Kell et al., 2018), their performance often diverges from biological auditory systems. This divergence manifests in their internal representational geometries and their vulnerability to adversarial attacks that do not impair biological hearing (Carlini & Wagner, 2018; Feather et al., 2023; Muzellec & Kar, 2026; Subramanian et al., 2019; Tuckute et al., 2023; Weerts et al., 2022). We addressed this discrepancy by reversing the order of the typical task optimization workflow: instead of assessing whether a category-optimized network predicts cortical responses, we evaluated whether our brain-optimized foundation model (ACNet) spontaneously developed categorical representations, and whether these representations were aligned with actual brain responses. To that end, we analyzed ACNet’s representations of the ESC-50 dataset (Piczak, 2015), an out-of-distribution corpus of environmental sounds not utilized during model training. To test the brain alignment of these representations, we also recorded neural responses to the same ESC-50 stimuli from a separate population of cortical neurons, independent of the ACNet training cohort.

A UMAP plot (McInnes et al., 2020) of the ACNet manifold embeddings revealed semantic clustering across most sound classes (Figures 3A and S1). Moreover, semantically related categories occupied adjacent regions in the projection space. For example, values in the first UMAP axis were low for *Animal* sounds but high for *Interior* sounds (Figure S1). To quantify how well the ACNet manifold can account for category decoding of neural populations in AC, we trained classifiers based on the spike counts in recorded neural data and on the manifold embeddings in ACNet (Figure 3B). We used time-averaged rates as classifier input since auditory cortical neurons carry substantial categorical information in their average rate (Bagur et al., 2025; Peng et al., 2024). To isolate the contribution of the network architecture itself from the effects of active training on neural activity, we trained two control classifiers with the following inputs: manifold embeddings of a model with ACNet architecture but shuffled linear weights, and the time-averaged stimulus spectrogram that provides the input to ACNet.

**Figure 3.**
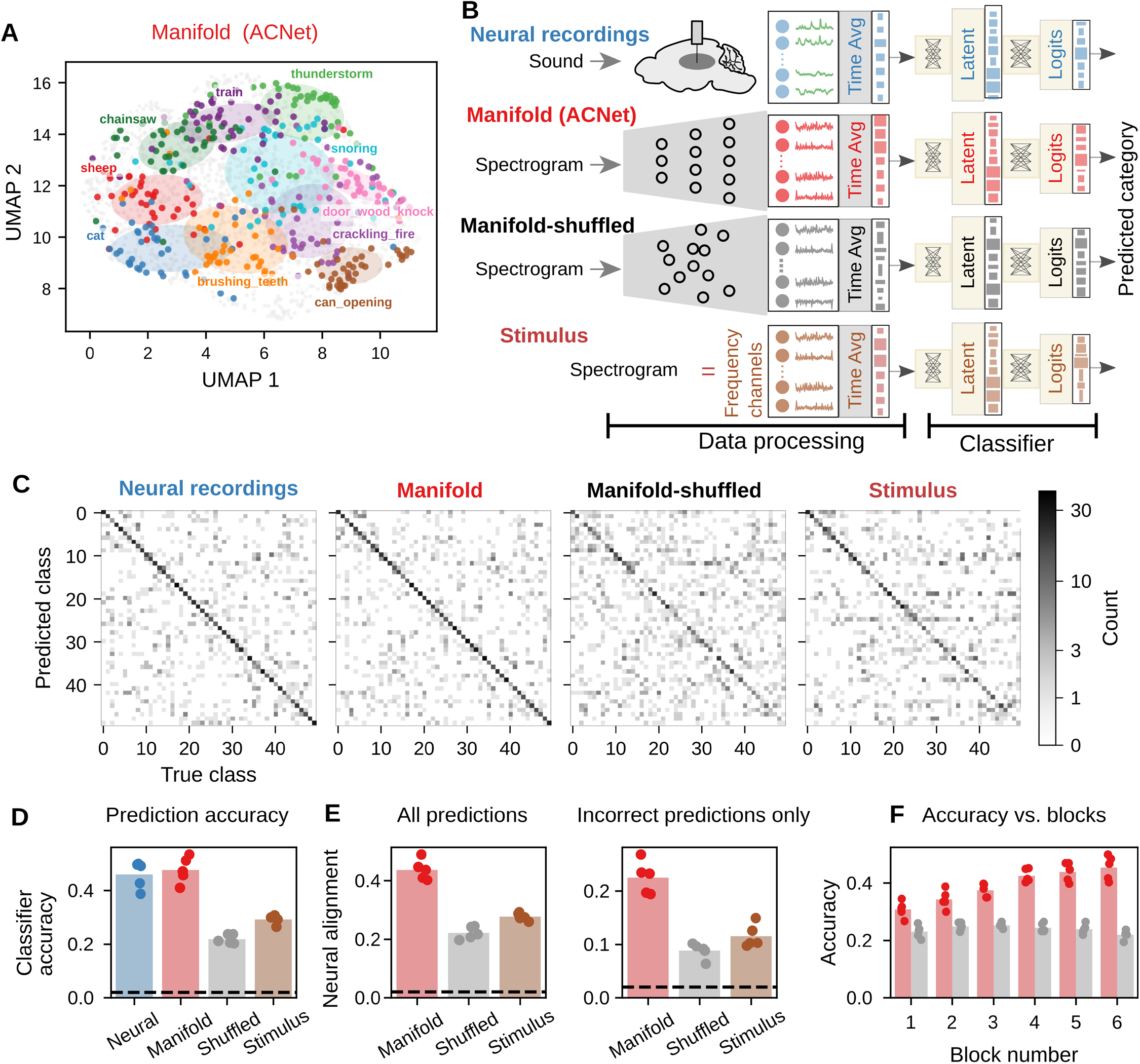
Manifold embeddings can be used to train neurally aligned sound-category classifiers. **(A)** UMAP plot of time-averaged ACNet manifold embeddings for ESC-50 stimuli. 10 exemplar sound categories are highlighted (colors) and stimuli from other categories are shown as gray dots. Ellipses demarcate one standard deviation along the major and minor axes for the highlighted categories. UMAP plot for all 50 categories is shown in Figure S1. **(B)** Schematic of classifiers trained using different inputs: neural (recorded PSTH), manifold (of ACNet), shuffled (after permuting the weights of ACNet in each layer), and stimulus spectrogram. All inputs were time averaged over the 0.5 sec stimulus sample. All classifiers had the same architecture (256 latent units and 50 output units). **(C)** Confusion matrices show the frequency of each predicted category versus true category for the four classifiers. See Figure S2 for confusion matrices for incorrect predictions only. **(D)** Mean prediction accuracy for the four classifiers. Dots represent five-fold jackknife estimates. Prediction accuracy (mean ± standard deviation per category) is as follows: Neural: 0.45 ± 0.16, Manifold: 0.47 ± 0.14, Shuffled: 0.20 ± 0.12, Stimulus: 0.31 ± 0.15. **(E)** Neural alignment, i.e., correlation coefficient between the confusion matrices for the neural data and the three model-based classifiers, considering all test stimuli (left) or incorrect predictions only (right). For all stimuli (Wilcoxon signed-rank test): Manifold > Shuffled: p= 5.3e-15, Manifold > Stimulus: p=1.6e-8. For incorrect stimuli: Manifold > Shuffled: p=7.7e-9, Manifold > Stimulus: p=9.0e-7. **(F)** Classification accuracy across convolutional blocks of ACNet (red) or weight-shuffled ACNet (gray). Accuracy increased significantly across blocks for ACNet (Spearman rank-order correlation: r=1.0, p<1e-16), but not for weight-shuffled ACNet (r=-0.37, p=0.47).

The neural- and manifold-based classifiers performed similarly (two-sided Wilcoxon signed-rank test, n = 50 categories, p = 0.21, n.s.), and they outperformed the shuffled manifold and stimulus-based classifiers (p < 3.2e-6, one-sided Wilcoxon signed-rank test, Bonferroni-corrected across all 4 pairwise tests; Figure 3D). While neural alignment should lead to similar overall accuracy between classifiers, it should also produce a similar pattern of errors, as reflected in the confusion matrix (Figure 3C). When we compared error patterns, the neural classifier showed significantly higher similarity to the manifold-based classifier than the shuffled-ACNet and stimulus-based classifiers (p < 9.0e-7, one-sided Wilcoxon signed-rank test, Bonferroni-corrected, Figures 3E and S2).

To test how this rate-based category representation emerges across successive convolutional blocks of ACNet, we trained a classifier based on each convolutional block’s embedding (Figure 3F). To determine whether this emergence simply reflected the network architecture (i.e., independent of whether it was trained to predict dynamic neural activity), we also trained a classifier based on a version of ACNet with shuffled weights. Classification accuracy increased along successive blocks of ACNet (Spearman rank-order correlation: r=1.0, p<1e-16) but not for shuffled ACNet (Spearman rank-order correlation: r=-0.37, p=0.47). These results demonstrate that accurate and neurally aligned categorical selectivity emerged in ACNet as a consequence of training to predict cortical population activity, rather than being a function of its structural network architecture or the spectral properties of the stimuli.

### Rate coding and sparse representation emerge across layers of ACNet

The analyses of response prediction accuracy (Figure 1) and category decoding (Figure 3) demonstrate that the multi-layer architecture of ACNet captures emergent sensory representations in AC. How does sound encoding change across successive layers of ACNet to produce these representations? Previous studies have shown that the dimensionality of evoked population activity increases and coding redundancy decreases along the auditory pathway (Atencio et al., 2012; Chechik et al., 2006; Gosselin et al., 2025). To test whether this result holds for ACNet, we obtained activations across successive model blocks in response to 100 training stimuli (∼33 min), which matches the typical fit dataset size per site.

To evaluate how coding redundancy changes across layers of ACNet, we applied PCA to the output of each convolutional block (Figures 4A and 4B). For each block, we fit a power-law function to its eigenspectrum, to measure variance explained per PC dimension. The exponent of this function quantifies the redundancy or efficiency of population coding, with higher values (shallower slopes) indicating increased efficiency (Stringer et al., 2019). The slope was systematically shallower along the ACNet hierarchy (Figure 4C), consistent with progressively higher encoding efficiency and greater independence between channels. These findings replicate previous findings that information redundancy is reduced along the auditory hierarchy (Chechik et al., 2006). To measure the dimensionality of the population code, we computed the participation ratio of the eigenspectrum for the output from each model block (Recanatesi et al., 2022). The participation ratio increased along the ACNet hierarchy (Figure 4D), consistent with increased dimensionality along the auditory pathway (Atencio et al., 2012; Gosselin et al., 2025). To verify that these results do not simply reflect the increasing number of units in deeper ACNet layers, we analyzed the activations of a control network with a flat architecture that had an equal number of units across all layers but an otherwise identical design. This flat-architecture control replicated the hierarchical dimensionality expansion of the main model (Figure S3). Because information cannot be created along successive feedforward operations (Cover, 1999), these results suggest a response sparsification, where information is distributed across an increasingly greater number of dimensions.

**Figure 4.**
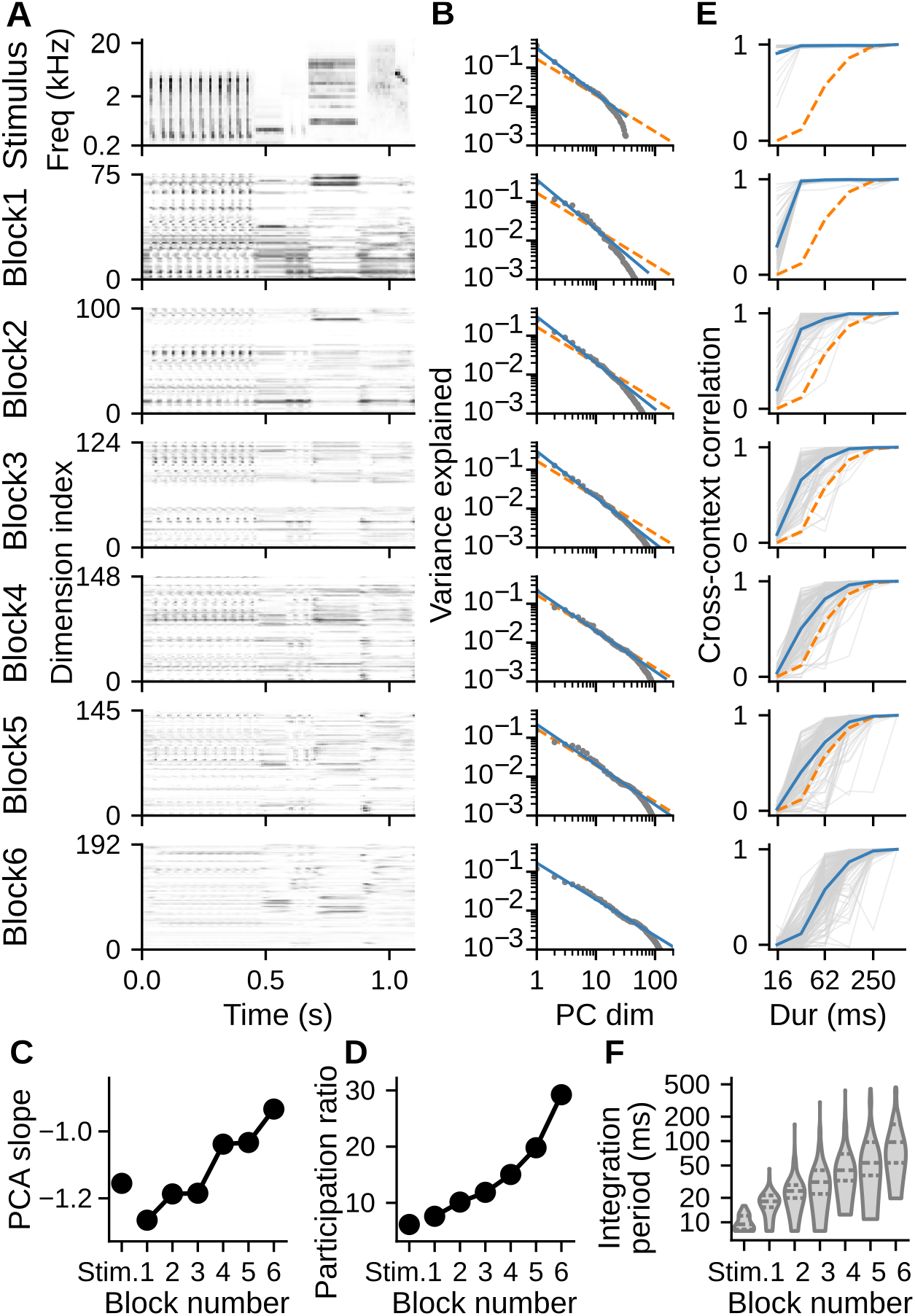
Emergence of joint rate and temporal coding in ACNet, concomitant with increased efficiency, dimensionality, sparsity, and diversity of integration time constants. **(A)** Example 1-s natural sound gammatone spectrogram (top) and corresponding model activations across convolutional blocks of ACNet. Decreased salience of temporal modulation across blocks highlights a temporal-to-rate code transformation. Block 6 is the encoding layer that defines the ACNet manifold. **(B)** Variance explained (gray) by principal components of activations for each block in A. Blue lines indicate linear fits (in log-log scale) for dimensions spanning 10**-**90% of the cumulative variance explained. Orange dashed lines indicate fits for the ACNet manifold (Block 6), which has a shallower slope. **(C)** Slopes of linear fits in B. Higher (i.e., less negative) values indicate slower decay in relative contributions across PC dimensions, consistent with increased efficiency (i.e., decreased redundancy). Slopes increased significantly across blocks (Spearman rank-order correlation = 1.0, p<1e-16). **(D)** Participation ratio, the effective dimensionality over which PC variance is spread, plotted per ACNet block, as in C. Participation ratio significantly increased across blocks (Spearman rank-order correlation = 1.0, p<1e-16). **(E)** Cross-context correlation (CCC) for individual channels in each block (thin gray lines), plotted against segment duration (log axis). Blue lines show population-averaged CCC. Dashed orange lines represent CCC for the ACNet manifold (Block 6). **(F)** Violin plots show distribution of temporal integration windows for all units in each block, measured as the width of a Gamma function fit to the CCC. Median temporal integration window increased across blocks (respectively, 9, 18, 28, 37, 52, 64, and 89 ms, Spearman rank-order correlation: r = 0.70, p < 1e-16), and the spread of integration windows increased (mean absolute deviation: Spearman rank-order correlation: r = 0.54, p < 1e-16), consistent with greater diversity of temporal integration across channels in later blocks.

How is the representation of sound transformed as it passes through ACNet? Previous studies have shown that temporal-to-rate coding conversion is ubiquitous along the neural auditory pathway (Joris et al., 2004; X. Wang et al., 2008), with the length of the temporal integration window also increasing along the auditory hierarchy (Asokan et al., 2021; Norman-Haignere et al., 2022). Inspection of activations in deeper layers shows some channels that faithfully track sound modulations, consistent with a temporal code, while others fluctuate more slowly, consistent with a rate code (Figure 4A). Thus, we considered how the balance of temporal and rate coding shifts across ACNet layers. To characterize temporal dynamics, we estimated the temporal integration window for units in each layer using the context invariance paradigm developed in (Sabat et al., 2025). Model activations were obtained for sequences of stimulus segments of different durations. Each segment was presented in two different sequences, thus occurring after two different preceding stimuli (contexts). The time point after which a unit responds similarly to stimuli in both contexts reflects its temporal integration period and was measured using cross-context correlation (CCC) of the unit activations (Figure 4E, see Methods). Integration window (*T*), the width of a Gamma distribution fit to the CCC (Figure S4), progressively increased along the hierarchy of ACNet. The gammatone representation had very short integration times (median *T* = 9 ms, close to gammatone time resolution), while the manifold layer had much longer context dependence (median *T* = 89 ms). Additionally, the distribution of *T* widened significantly across layers, as indicated by increasing mean absolute deviation (Figure 4F). This wide distribution of time constants could support parallel rate and temporal codes in deeper layers, as representations become higher-dimensional and more independent.

### ACNet captures both temporal and rate aspects of neural coding

Neural activity patterns in the auditory cortex are typically characterized by two distinct coding regimes: a temporal pattern that is synchronized to the stimulus envelope and a spike rate that encodes stimulus features asynchronously (Lu et al., 2001). We tested whether ACNet could account for these fundamental modes of cortical coding. We simulated responses to click train stimuli with variable inter-click interval (ICI), classically used to distinguish temporal and rate codes (Figure 5A). The key response properties that define these modes are whether responses are phase locked to the stimulus each cycle (the *synchronized* group) or whether responses vary in the spike count but not spike timing (the *nonsynchronized* group). We classified responses according to these criteria, assigning responses that matched both criteria or neither to *dual* and *none* groups, respectively (Figure 5C, see Methods for details about group assignment).

**Figure 5.**
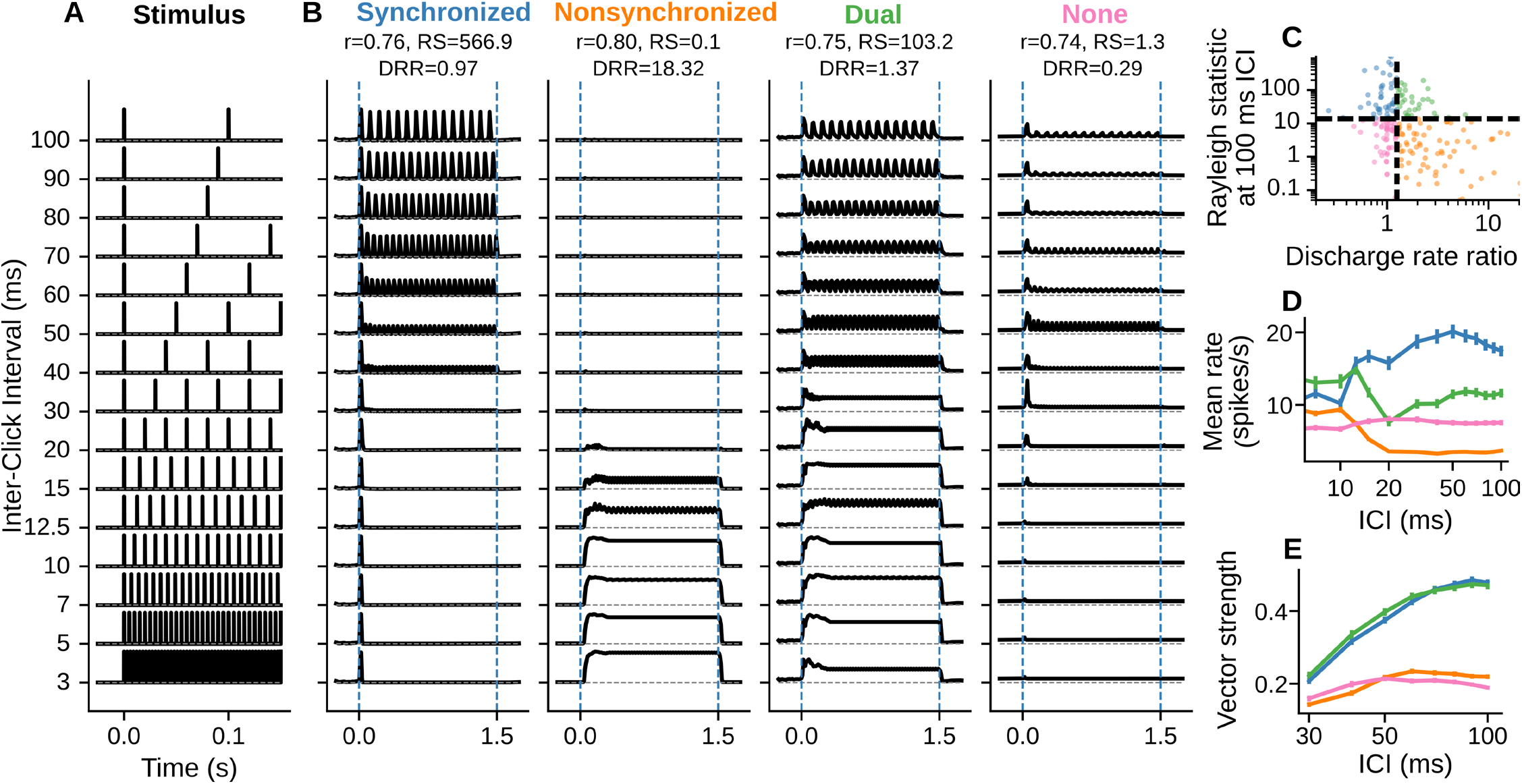
ACNet captures both rate and temporal coding of click-train stimuli. **(A)** Click train stimuli spanning fast (3 ms) to slow (100 ms) inter-click intervals (ICIs) were used to obtain simulated neural PSTH responses from ACNet (1.5 s duration, 150-ms segment plotted). **(B)** Simulated neural responses for four example neurons typical of categories defined by their alignment with sound envelope: synchronized (blue), nonsynchronized (orange), dual (green), and none (pink). Text indicates ACNet prediction accuracy for validation stimuli (r), Rayleigh statistic (RS, a measure of temporal coding), and discharge rate ratio (DRR, a measure of rate coding) for each neuron. **(C)** Scatter plot of DRR versus RS at 100 ms ICI for all simulated responses shows neurons falling into four categories, defined by boundaries at DRR=1.25 and RS=13.8. **(D)** Mean time-averaged firing rate for the four groups. Error bars show 1 SEM. **(E)** Mean vector strength for the four groups. Error bars show 1 SEM. Colors in **C-E** indicate category, as in B.

As expected, *synchronized* neurons showed higher firing rate and stronger stimulus-synchronized activity at higher ICIs (> 30 ms, Figure 5E) and weaker firing rate at shorter ICIs (Figure 5D). These responses not only included stimulus-excited synchronization, but also phase-locked disinhibition for neurons suppressed at faster click rates (Figure S5A). Nonsynchronized neurons encoded shorter ICIs (<20 ms) in their firing rates, but they showed little modulation in firing rate at longer ICIs or vector strength across ICIs (Figures 5D and 5E). This category included neurons that completely discarded temporal patterns (e.g., a step response) or show brief onset or offset transients, or their combination (Figure S5B). Dual neurons multiplexed both strategies, employing rate coding at shorter ICIs and switching to a temporal code at longer ICIs (Figures 5B–5E). These neurons also showed variable response patterns at stimulus onset and offset, as well as enhanced or suppressed phase-locked responses during the sustained period (Figure S5C). Responses of *none* neurons did not meet these criteria; the population response showed little tuning to fast or slow ICIs (Figures 5B, 5D, and 5E). This group contained several subcategories, including onset-only responses without slow synchrony, band-pass tuning in temporal or rate responses, or flat rate profiles across ICIs (Figure S5D).

Natural sounds often contain regularities like those in click trains, and thus we could test for similar patterns of temporal coding in the recorded neural data. We analyzed responses to two natural sound segments in the validation stimuli, drums and chirps, with temporal structures analogous to fast and slow click trains, respectively. These responses exhibited the same range of profiles as ACNet, including stimulus-excited or stimulus-inhibited synchronized activity, and onset, offset, and sustained nonsynchronized activity patterns (Figure S6). Thus, the parallel temporal and rate-based coding observed in ACNet can be observed directly in responses to natural sound stimuli.

### Tuning properties of the auditory cortical manifold

What tuning properties does ACNet learn that allow it to account for representations observed across the entire neural population? To address this question, we focused on the *encoding manifold*, which is the shared representational bottleneck common to all neurons in our dataset. Since the mapping between manifold activation and neural PSTH response is a memoryless transformation (i.e., a linear combination of manifold outputs followed by a static nonlinearity), the manifold consists of a set of basis functions trained to best predict instantaneous neural population activity.

We observed substantial heterogeneity in sensory tuning across manifold dimensions. Activations for a sequence of synthetic stimuli, including pure tones, sinusoidally amplitude-modulated (SAM) tones, SAM noises, upward and downward frequency sweeps, and click trains (Figure 6A) showed diverse response patterns across manifold dimensions (Figure 6B). Crucially, the simulated profiles revealed a clear division in temporal processing: some dimensions acted as fast temporal filters, phase-locking precisely to individual sweeps and clicks, whereas others behaved as slow rate-integrators, tracking overall envelope energy with minimal temporal locking.

**Figure 6.**
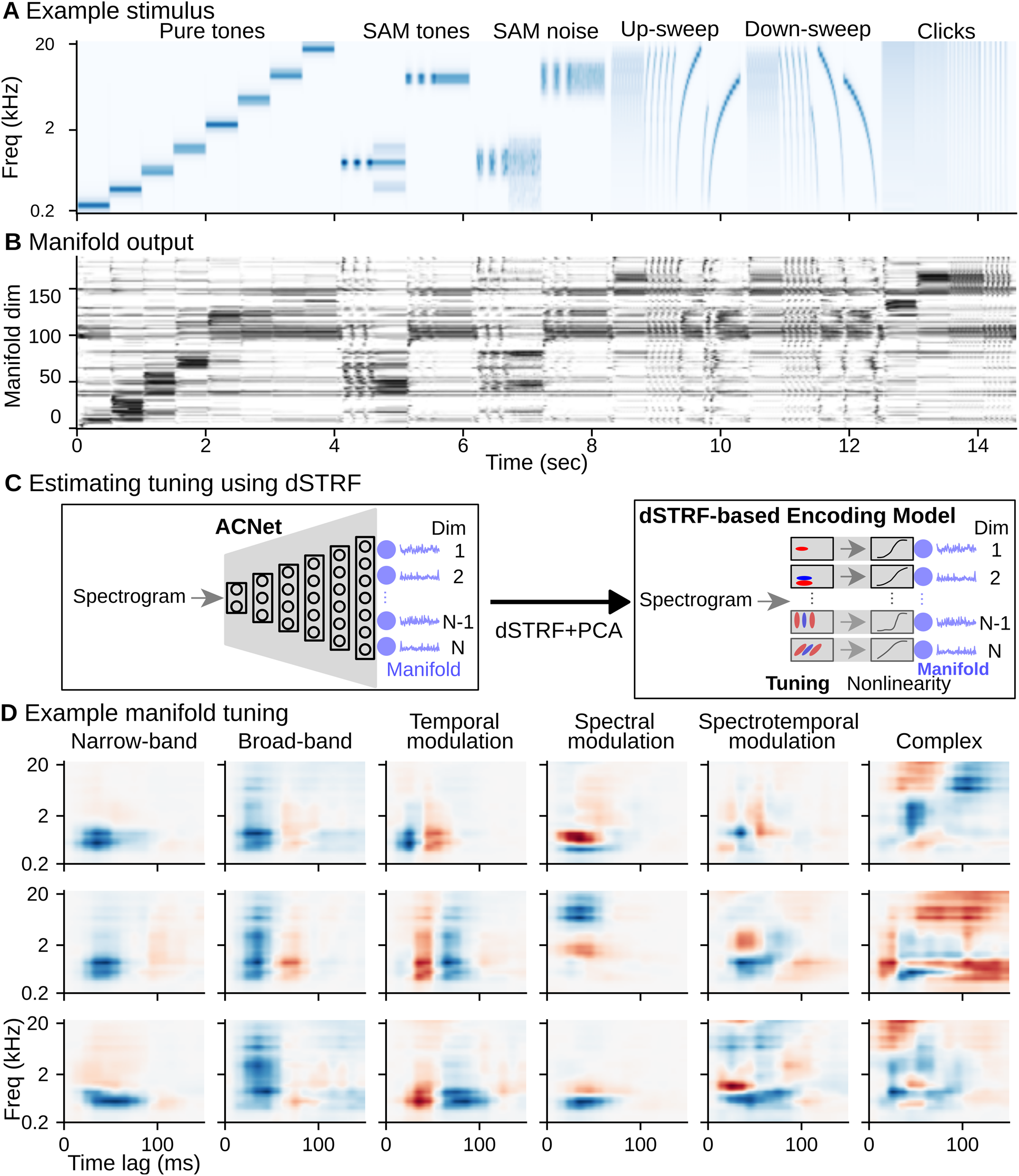
Manifold represents a set of basis functions, capturing a diversity of tuning properties. **(A)** Spectrogram of synthetic stimulus battery presented to ACNet, which included pure tones, SAM tones, SAM noise, upward and downward sweeps, and click trains. **(B)** Heatmap plots of activation of individual manifold dimensions by the synthetic stimulus sequence in A. **(C)** The spectrotemporal receptive field (STRF) of each manifold dimension was measured using dynamic spectrotemporal receptive field (dSTRF) analysis, which uses the gradients of the response with respect to the stimulus to identify the dominant STRF affecting each manifold dimension. A subset of the natural stimuli used for ACNet training was used to estimate dSTRFs. **(D)** The STRFs (first principal component of dSTRFs) of example manifold dimensions highlight distinct patterns of spectrotemporal tuning, denoted by each column.

While the simulated responses highlight the diversity of manifold basis functions, these responses do not reveal the spectrotemporal patterns that evoke these activations. To that end, we used dynamic spectrotemporal receptive field (dSTRF) analysis (Wingert et al., 2026) to characterize the STRF of each manifold dimension. To ensure that dSTRF tuning reflects sound processing for natural stimuli, we used a subset of the natural sound sequences that was used for ACNet training (see Methods). To obtain STRFs, we measured the gradient of each manifold dimension output with respect to the preceding input stimulus, identifying features that influence the manifold output at each time point. We then performed dimensionality reduction (PCA) and identified the dominant tuning feature, or STRF, as the highest variance PC for each manifold dimension (schematized in Figure 6C). The range of tuning for different manifold dimensions (Figure 6D) demonstrated a diversity of spectrotemporal tuning that underlies the selective responses to synthetic stimuli in Figure 6B. Tuning of these dimensions spanned narrow to broadband frequency tuning, temporal/spectral/spectrotemporal modulation tuning, and complex tuning. We also observed similar diversity in the frequency response area (FRA) for different dimensions (Figure S7), which is classically used to characterize joint frequency-intensity tuning. Thus, the manifold basis functions support a rich set of time-varying activations to simultaneously support precise temporal patterns and robust rate-based representations across the cortical population.

### AC manifold reveals distinct functional properties across cell types

Previous studies have shown that tuning similarity between neurons can vary across cortical regions and cell types (Atencio & Schreiner, 2010; Bizley et al., 2005; Liu & Kanold, 2021; Maor et al., 2016; Polley et al., 2007; Sabat et al., 2025), although the extent and consistency of these differences remain unclear (Landemard et al., 2021). Therefore, we next asked whether the manifold can reveal differences in tuning properties between neural populations defined by their anatomy (Figure 7A). We took advantage of the fact that the manifold represents a single basis set for activity of the entire neural population. Thus, each neuron has a distinct loading vector, characterizing the influence of each manifold dimension on its activity. To compare the functional properties of two neurons, we computed their manifold loading similarity (MLS), defined by their cosine similarity, which ranged from -1 (anticorrelated) to +1 (perfectly correlated). We tested three specific hypotheses: 1) whether anatomically nearby neurons (i.e., within the same cortical column) have more similar tuning (higher MLS) than distant neurons (i.e., in different recording sites), 2) whether MLS varies between primary (A1) and secondary (PEG) auditory cortices, and 3) whether the ACNet manifold is truly generalizable, i.e., if MLS for neurons in different animals is similar to that of distant neurons in the same animal.

**Figure 7.**
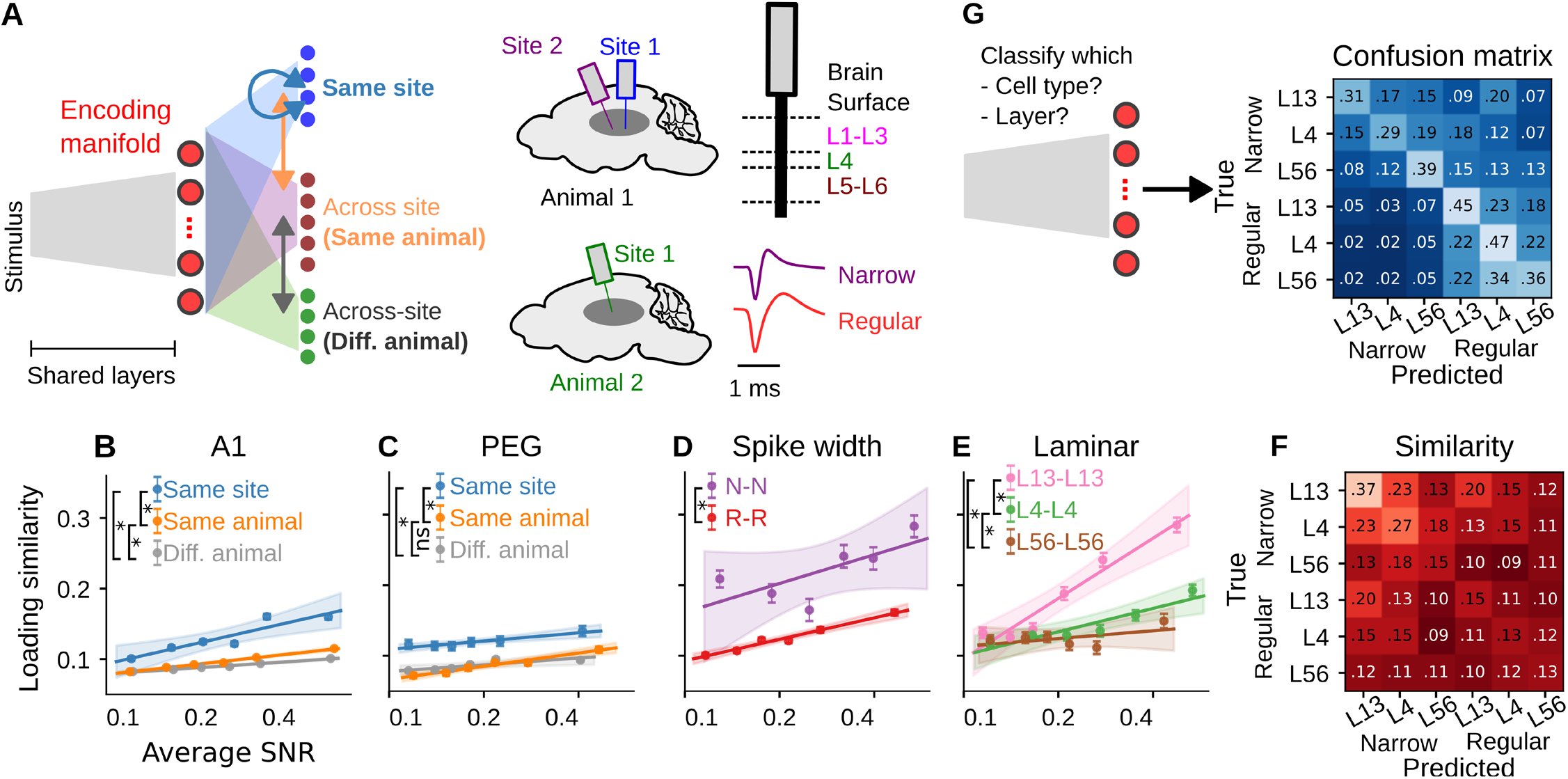
Manifold-to-neural loading describes tuning similarity across anatomical regions and cell types. **(A)** Schematic of manifold-to-neural loading comparisons. **(B)** Manifold loading similarity (MLS) for pairs of cells within the same recording site (blue, *Same site*, count = 15,066 pairs), different sites in the same animal (orange, *Same animal*, count = 190,848), and in different animals (gray, *Diff. animal*, count = 341,197) for A1. The abscissa (x-value) for each pair is the geometric mean of response signal-to-noise ratio (SNR) for the two cells. The ordinate for each group corresponds to residual MLS for that group (Equation 2). Points show binned means ± SEM; shading shows 95% confidence intervals. Parenthetic numbers correspond to the number of pairs in each condition. Detailed statistics are reported in Supplemental Table S2. **(C)** Same as B, but for data from secondary field PEG (count: Same site = 4,257, *Same animal* = 6,720, Diff. animal = 25,642). A1 and PEG were not significantly different (PEG-A1: *β* =-0.0056, 95% CI [-0.0258, 0.0193]; Supplemental Table S2); therefore, data are pooled across A1 and PEG in remaining panels. **(D)** MLS for pairs of neurons within a site with narrow (*N-N*, putative inhibitory neurons, count = 1,254) or regular (*R-R*, putative excitatory neurons, count = 12,178) spike widths. MLS for *N-N* was significantly greater than that for R-R (*β* =-0.0796, 95% CI [- 0.0921, -0.0671], Supplemental Table S4). **(E)** MLS for pairs of neurons in the same layer within a site: supra-granular (*L13-L13*; count = 2,779), granular (*L4-L4*; count = 3,094), and infra-granular (*L5C-L5C*; count = 2,648). MLS decreased from supragranular (L13) to infragranular (L56) layers (L13 vs. L4: *β* =0.0510, 95% CI [0.0414, 0.0604]; L4 vs. L56: *β* =0.0133, 95% CI [0.0016, 0.0248]; Supplemental Table S4). **(F)** Marginal residual MLS for all pairs within a site, indicating loading similarity after accounting for random effects. MLS across sites is shown in Figure S8. **(G)** Confusion matrix for joint cell-type and lamina classification from MLS (mean macro-F1 = 0.38; chance = 0.15 ± 0.01; permutation test, p < 0.001, 1000 label permutations, z = 18.6). Accuracy measures are for held-out test recording sites (five-fold site jackknife)

To address these hypotheses, we constructed a Bayesian linear mixed-effects model (see *Methods*) with these regions and site-connectivity as predictors and MLS as the outcome (Figures 7B and 7C). Since response SNR is correlated with tuning similarity between pairs of neurons (Wingert et al., 2026), it was included as a predictor. Model response prediction accuracy was also included as a predictor to control for the possibility that poorly predicted neurons may map onto the same noisy manifold dimensions. Site identity of both neurons was treated as a random effect. MLS was significantly higher for pairs of neurons within-site than across sites, confirming the role of anatomical proximity in shaping sensory tuning (*β* = 0.0391, 95% CI [0.0351, 0.0432]). MLS was not significantly different between A1 and PEG (*β* = -0.0056, 95% CI [-0.0258, 0.0193]) in this model. However, this lack of difference when considering only pairs of neurons from the same area does not preclude the possibility that A1 and PEG have different manifold loadings, but similar within-area MLS. We tested this possibility by constructing another Bayesian model including neuron pairs from different areas and confirmed that A1 and PEG were indeed not statistically different (Supplemental Table S3). MLS for within-animal (different sites) and across-animal was not different for PEG (*β* = -0.0001, 95% CI [-0.0082, 0.0079]) and marginally different for A1 (*β* = -0.0074, 95% CI [- 0.0090, -0.0057]; Supplemental Table S2). Overall, these results suggest that the manifold captures transformations of auditory stimuli that are anatomically locally clustered, yet globally generalizable across subjects, demonstrating that the latent computational space is highly conserved.

Previous studies have also shown differences in tuning properties between cell types and layers in the cortical microcircuit (Atencio & Schreiner, 2010; Moerel et al., 2019; Montes-Lourido et al., 2021; Williamson & Polley, 2019; Winkowski & Kanold, 2013). Thus, we asked whether MLS, which measures tuning similarity, can reveal similarity and differences between these cell types. Specifically, we asked whether MLS was different across cell types (putative excitatory [regular spike width] and putative inhibitory [narrow spike width]) and across cortical laminae (supra-granular [L1-L3], granular [L4], and infra-granular [L5-L6]). To maximize statistical power for microcircuit-level analyses, we pooled data across A1 and PEG because MLS was similar between these fields (Supplemental Tables S2 and S3). To test MLS similarity at both local and global anatomical scales, we constructed two separate Bayesian mixed-effects models for MLS *within-site* (Figures 7D–7F; Supplemental Table S4) and *across-site* (pooled across same and different animals, Figure S8; Supplemental Table S5). Within-site, MLS for narrow-spiking neurons was significantly higher than for regular-spiking neurons (regular-to-narrow contrast: *β* =-0.0796, 95% CI [-0.0921, -0.0671]; Figure 7D), consistent with higher tuning specificity (thus lower generalizability) of regular-spiking neurons (Liu & Kanold, 2021; Maor et al., 2016). Across laminae, MLS decreased from superficial to deeper layers, i.e., L13 > L4 > L56 (Figure 7E), consistent with previous reports (Atencio & Schreiner, 2010; Wingert et al., 2026). Trends of these effects were similar for both within- and across-site conditions, though the effects were weaker for across-site than within-site comparisons (Figures 7D–7F and S8; Supplemental Tables S4 and S5).

While MLS reveals average tuning similarity between groups of neurons, e.g., across cortical laminae, it does not address whether these groups are identifiable based on their manifold-to-neural loading. For example, a group of neurons that primarily receive orthogonal projections (loadings) from a few manifold dimensions will have low MLS within group, but it will remain identifiable if those dimensions are not used by other groups. Therefore, we next asked to what extent these manifold-to-neural loadings can be used to classify different cell types and cortical laminae, as has been shown in a foundation model of vision (E. Y. Wang et al., 2025). We trained a simple two-layer perceptron to classify both cell type and cortical lamina simultaneously (trained and tested on non-overlapping sites). The overall test classification macro-F1 score was 0.38, which was significantly greater than chance (0.15; Figure 7G). Thus, in addition to accounting for differences in tuning between cortical sites and layers, the ACNet manifold captures generalizable sensory transformations that distinguish different cell types, regardless of cortical site.

## Discussion

We have developed ACNet, a foundation model for sound encoding by auditory cortex (AC) that generalizes across individuals and large neural populations within individuals. ACNet captures several key functional properties of cortex, including temporal-to-rate transformations along the auditory hierarchy, concomitant with increased dimensionality, sparseness, and diversity in temporal integration in layer representations (Figure 4). These transformations enable neural responses to multiplex information in both their timing and rate – a fundamental dichotomy in neural coding. Dimensionality analysis revealed that AC populations represent sensory stimuli with the same near-optimal efficiency as is observed in the visual cortex (cvPCA slope ≈ -1, Figure 1E) (Stringer et al., 2019), suggesting that encoding optimality is a universal principle in sensory cortex. In contrast to previous models focusing on specific aspects of neural coding (Bendor, 2015; Bondanelli et al., 2021; Chambers et al., 2019; David & Shamma, 2013; Fishbach et al., 2001; Kudela et al., 2018; Mesgarani et al., 2014; Parida et al., 2022; Rançon et al., 2024), ACNet unifies response properties spanning phase-locked, onset, offset, stimulus-excited and stimulus-suppressed activity (Figures S5 and S6) with state-of-the-art prediction accuracy (Figure 1).

### Generalizability of the ACNet manifold

An architecture combining shared encoding layers and recording site-specific decoding layers permits ACNet to account for the activity of thousands of neurons in multiple animals with a relatively low-dimensional encoding manifold. Previous studies have reported across-subject similarities in evoked neural activity in several sensory modalities (P.-H. (Cameron) Chen et al., 2015; Güçlü & van Gerven, 2017; Haxby et al., 2011; Sabesan et al., 2023; Schneider et al., 2023; Yang et al., 2026); however, this work has typically used data that lack the temporal or anatomical specificity of single neurons in cortex. Thus, our results establish a baseline representational similarity between individuals at the single-neuron level in AC. Despite the large diversity of encoding properties across neurons recorded from different individuals, our encoding model approach highlights how the underlying geometry of these representations is shared at the level of the encoding manifold.

Whereas the ACNet encoder characterized the dominant sensory transformations relevant to population neural activity, the ACNet decoder could differentiate functionally distinct cell types using loadings in its decoder. Within the decoder, the manifold-to-neural loadings were informative about functional groups of cells (Figure 7), similar to findings for a foundation model of visual cortex (E. Y. Wang et al., 2025). For example, narrow-spiking neurons showed more similar loadings than regular-spiking neurons, consistent with the higher tuning diversity of regular-spiking neurons than narrow-spiking neurons (Liu & Kanold, 2021; Maor et al., 2016). Similarly, within-class similarity decreased from supra-granular to infra-granular layers, specifically for the narrow-spiking neurons (Figures 7F and S8C). These findings are consistent with previous reports (Atencio & Schreiner, 2010; Wingert et al., 2026) and highlight how sensory tuning varies across laminae within a cortical column. Our data did not show a significant difference between A1 and PEG, which could either reflect smaller differences in the ferret AC or that these differences emerge for sound mixtures or task demands (Atiani et al., 2014; Landemard et al., 2025). Overall, these results suggest that there exists a low-dimensional generalizable manifold that captures canonical sensory transformations in the cortex and facilitates the heterogeneous tuning observed across large neural populations and across different subjects.

### Manifold accounts for neural representations of sound category

A critical task of the auditory system is to categorize sounds. Modern audio-based classifiers are typically trained on a vast amount of data, enabling these models to achieve classification performance on par with or beyond that of humans (K. Chen et al., 2022; S. Chen et al., 2022; Gong et al., 2021; Kong et al., 2020; Tokozume et al., 2018). These models can also explain some aspects of human hearing, e.g., the effects of background noise and spectrotemporal filtering on categorical perception (Alavilli & McDermott, 2026; Kell et al., 2018). Despite their successes, these models can still underperform in some listening conditions (e.g., temporal masking), show categorical invariances and vulnerability to adversarial attack that are not human-like, and deviate in their internal representations from the brain (Adolfi et al., 2023; Carlini & Wagner, 2018; Feather et al., 2023; Muzellec & Kar, 2026; Subramanian et al., 2019; Tuckute et al., 2023; Weerts et al., 2022). Additionally, while these models have been shown to predict brain responses to various degrees, the brain data used typically lack the temporal and/or anatomical specificity of the auditory system. Here, we establish an alternative and more direct approach to build a neurally aligned classifier. Using the ACNet manifold as the front-end to the classifier ensures that the information into the classifier follows biological principles based on its training to predict neural activity at fast (10-ms) temporal resolution. Critically, this training allowed ACNet to build category selectivity for an out-of-distribution stimulus dataset (ESC-50, Figure 3), suggesting that categorical selectivity is an organizational principle of hierarchical processing in the auditory system, similar to other sensory modalities (Gottfried, 2010; Güçlü & Gerven, 2015; Li & DiCarlo, 2012). The ACNet-based classifier replicated the confusion pattern in an independent neural dataset better than the stimulus-based classifier. These results show that brain-based models, like ACNet, can be used for downstream applications. These models can also address several outstanding questions, such as the role of temporal-vs-rate information in category decoding (Bagur et al., 2025; Engineer et al., 2008; Kayser et al., 2010; Malone et al., 2010; Mesgarani et al., 2008; Walker et al., 2008) and whether these neurally based task-optimized models can reduce the divergence between artificial and biological neural networks.

### A spectrotemporal basis set for cortical sound representation

The shared encoding layers of ACNet provide a relatively compact, low-dimensional characterization of sound feature representation in AC. Spectrotemporal receptive fields (STRFs) measured by subspace analysis of ACNet manifold dimensions revealed a diversity of spectrotemporal tuning (Figure 6). These neurally derived transformations provide a more complete description of cortical function compared to previously hand-engineered approximations (e.g., a Gabor filter bank, Chi et al., 2005; Jones & Palmer, 1987). The overall tuning space of the manifold will be even more complex and diverse, as higher order dSTRF PCs add nonlinear interactions that increase the complexity and sparsity of these responses (Wingert et al., 2026). These hierarchical transformations allow a temporal-to-rate code transformation, supported by longer and more diverse temporal integration windows along the layers of ACNet, similar to that observed in the biological auditory system (Joris et al., 2004; X. Wang et al., 2008). Overall, the encoder-decoder stages of ACNet open several avenues of future research, e.g., comparing internal representations of ACNet and other task-optimized models and their ability to predict auditory perception, how manifold-based tuning varies across more precisely defined cell types (e.g., classes of inhibitory neurons), characterizing how the manifold is affected in various forms of sensorineural hearing loss, and building neurally based audio applications that replicate the invariant representations in the brain to acoustic degradations. More generally, ACNet can be integrated into more general-purpose foundation models of the brain, which incorporate multiple modalities and tasks into a single model (Azabou et al., 2023, 2025; d’Ascoli et al., 2026; Ortega Caro et al., 2024; Willeke et al., 2026; Zhang et al., 2025).

### Limitations and future work

While ACNet achieved prediction accuracy on par with current state-of-the-art CNNs, a significant portion (40-45%) of response variance remains unexplained. This difference could partly be due to data limitations for each neuron (Mazzaschi et al., 2025; Pennington & David, 2023; E. Y. Wang et al., 2025; Willeke et al., 2026) and the need for model architectures that support richer input representations than are captured by ACNet (e.g., finer spectrotemporal resolution (Drakopoulos et al., 2025)). Our work with ACNet establishes that an encoding model for cortical single-unit population activity can be used to perform a downstream task, such as classifying sounds. Future studies comparing ACNet-based models with a wide range of task-optimized (non-neural) models may help explain the existing divergence in behavioral (measured by confusion patterns) and representational (measured by similarity to brain activations) aspects of object classification (Carlini & Wagner, 2018; S. Chen et al., 2022; Gong et al., 2021; Kong et al., 2020; Tuckute et al., 2023; Weerts et al., 2022).

## Acknowledgments

This work was supported by the National Institutes of Health BRAIN Initiative grant R01EB028155 (SVD) and National Institute on Deafness and Other Communication Disorders grants K99DC022330 (SP) and R01DC014950 (SVD).

## Model availability

The trained ACNet model is available at https://github.com/sbp894/ACNet.

## Methods

All experimental procedures were approved by the Oregon Health & Science University Institutional Animal Care and Use Committee (IACUC) and conformed to the standards of the Association for Assessment and Accreditation of Laboratory Animal Care (AAALAC) and the United States Department of Agriculture (USDA). Five neutered young adult ferrets (*Mustela putorius furo*, four males, one female) were obtained from Marshall Farms at 6-8 months old. Four of these ferrets (one female) were used to train ACNet, and the remaining ferret was used to collect the ESC-50 dataset. To allow for semi-chronic, head-fixed neurophysiology recordings, each animal was surgically implanted with headposts (Hamersky et al., 2025). Anesthesia was induced with Ketamine (10 mg/kg) and Midazolam (0.3 mg/kg) and maintained with Isoflurane (1-3%). In a sterile setting, an incision was made at midline, followed by tissue clearance over the skull. Next, two stainless-steel head posts were cemented along the midline using UV-light-cured bone cement (Charisma, Kulzer). The implant was reinforced with 8-10 self-tapping set screws mounted in the skull. The final shape of the implant was built by layers of dental cement. For the duration of experiments, the cranial implant margin was regularly cleaned with dilute Chlorhexidine or Betadine, flushed with sterile saline, and followed by sterile bandaging. After the recovery period of two to three weeks, animals were gradually habituated to head fixation.

### Stimuli

#### Sound presentation

Sound files were processed through an analog-to-digital converter (DAC, National Instruments), amplified (Crown), and presented via a free-field speaker (Manger) placed 80 cm from the listening animal’s head at 0° elevation and 30° azimuth contralateral to the recording hemisphere. Stimuli were presented at 44 kHz sampling rate and controlled using custom MATLAB software (https://bitbucket.org/lbhb/baphy) or equivalent Python software (https://github.com/LBHB/psiexperiment). All experiments took place inside a sound-isolating chamber (Professional Model, GretchKen) with a custom double-wall insert.

#### Model training

Natural sound sequences were used to train encoding models to predict population neural activity from the input stimulus spectrogram and have been described previously (Wingert et al., 2026). Each sequence was 17.79 sec in duration and consisted of 124 segments (64 50-ms segments, 32 110-ms segments, 16 190-ms segments, 8 420-ms segments, 4 780-ms segments) and five silence segments (one per duration). The pseudo-logarithmic distribution of segment durations minimized harmonic ringing in neural responses. Constituent segments were from one of two large corpora: Audio Set (Gemmeke et al., 2017) or Pro Sound Effects (PSE, Core 3 Complete). Segment sound levels spanned a 20-dB range uniformly and were combined with crossfading (10-ms Hanning window) to minimize transition artifacts. These sequence stimuli for model fitting were played once per recording site, with each sequence presented to either one (n = 23), two (n = 28), or three (n = 18) sites.

#### Model testing

Stimuli for model testing included six sequences, four of which were constructed similar to those described for model training. Two additional 18-s sequences consisted of 12 1-s segments of natural and synthetic sounds, which had been tested in previous experiments from our laboratory (Pennington & David, 2023). These segments were separated by 500 ms of silence. The synthetic segments included tones and noises, and natural segments consisted of vocalizations, music, and environmental sounds. Test sequences were repeated between 10 and 20 times to produce a reliable peristimulus time histogram to test model performance. Training and test stimuli were presented between 55- and 65-dB SPL RMS level.

#### Auditory categorization

Categorical selectivity of the encoding manifold and neural responses was assessed with a dataset of environmental sounds (ESC-50; Piczak, 2015). These stimuli comprised 50 categories (40 exemplars per category, 2000 total exemplars). To make the duration of neurophysiological recordings with these stimuli tractable, we used a truncated version of this dataset by dividing each 5-s exemplar into 10 segments of 500 ms each and selecting the loudest segment for presentation. Exemplars were interleaved with 100-ms silence between them and were repeated 10 times. Stimuli were presented at 55 dB SPL. The same stimuli were used for all classifiers (Figure 3).

### Neurophysiology

We recorded extracellular neural activity from awake, head-fixed ferrets passively listening to sounds. For each recording, we performed a small (0.5-1 mm diameter) craniotomy over the AC, guided by established tonotopic and anatomical landmarks (Bizley et al., 2005). Response properties were further analyzed to verify cortical subfields (Bizley et al., 2005). Primary AC (A1) was identified by tonotopic responses across multiple penetrations with short latency. The secondary AC field PEG was identified as ventrolateral to A1, with the boundary defined by a reversal of the tonotopic gradient at low frequencies.

Data were recorded either with an acute insertion of a 64-channel silicon electrode array (1.05-mm long; one animal) or a semi-chronic implantation of Neuropixels short NHP probes (IMEC, 384/960 selectable channels). Penetrations were perpendicular to the surface of the skull. Probe tips were either sharpened in-house or bought pre-sharpened to facilitate penetration through dura. In the Neuropixels preparation, probes were attached to a manual microdrive and mounted on a detachable base (Vöröslakos et al., 2021). For each recording, the microdrive and base were tightened, and the base was secured on the skull using dental cement. A silicone sealant was used to stabilize the microelectrode shank. The probe was allowed to settle for at least 18 hours following implant to improve recording stability. Once the recordings were completed, the microdrive-probe assembly was detached from the base for explanting. The electrodes were sterilized with 2% Glutaraldehyde (2-3 hours) followed by a saline rinse (30 cc) before implant and with 0.25% Trypsin (1-2 hours) followed by rinsing with deionized water after explant.

Electrophysiological signals were amplified (RHD 128-channel headstage, Intan Technologies, or Neuropixels headstage, IMEC), digitized at 30 kHz (Open Ephys), and saved to disk for further analysis. Spike sorting was performed offline with Kilosort2 (https://github.com/MouseLand/Kilosort2) and sorting results were manually curated in phy (https://github.com/cortex-lab/phy). Units with low (<5%) contamination parameter in the inter-spike histogram, well-separated feature cluster, and stable feature and amplitude profiles were classified as single units. Only responses from single units were included in this study.

#### Cortical layer classification

To identify cortical layer boundaries (1-3, supragranular; 4, granular; and 5-6, infragranular) in each recording, we used current source density (CSD) analysis (Kajikawa & Schroeder, 2011). Trial-averaged (60 trials) responses to a narrowband noise stimulus (2-6 octaves centered at the best population frequency measured using tone pips) were used for this analysis. We constructed the local field potential (LFP) signal by low-pass filtering the raw signal below 250 Hz (zero-phase, effectively fourth-order Butterworth filter). The second-order derivative of the LFP was estimated as the CSD.

Layer boundaries were estimated manually by visual inspection using a GUI (https://github.com/LBHB/laminar_tools). Average noise-evoked responses were sorted by depth and inspected for patterns of CSD source and sink evoked by the noise stimulus. These patterns were matched with the average CSD patterns evoked for AC previously reported across multiple species (Mendoza-Halliday et al., 2024; Schaefer et al., 2015). For each curated single unit, the layer was assigned based on the depth of the channel with the largest spike template amplitude for that unit.

#### Spike width classification

We classified neurons as narrow or regular spiking based on the width (time between depolarization trough and hyperpolarization peak) of the average action potential waveform (Trainito et al., 2019; Wingert et al., 2026). The width threshold for categories was 0.35 ms for the 64-channel probes and 0.375 ms for the Neuropixels probes, shifted to accommodate the distinct filtering properties each system applied to neurophysiological traces. We have previously shown that GABAergic inhibitory neurons in the ferret AC correspond to the narrow spiking group (López Espejo & David, 2024).

### Encoding model architecture

We tested several encoding model architectures, including previously published convolutional neural network (CNN) architectures that yield state-of-the-art performance for single recording sites. The architecture differed from our previous CNN models (Wingert et al., 2026) in two key ways. First, the models incorporated residual convolutional blocks, which reduce the vanishing gradient problem in deep neural networks (He et al., 2016). In each residual convolutional block, the main branch consisted of temporal convolution (finite impulse response filtering), followed by a linear (dense) layer, whereas the residual branch was a linear layer. The sum of the two branches went through a nonlinearity (ReLU). The scale of the main branch (relative to the residual branch) was learnable and was initialized to one. Second, model training used gradient accumulation across multiple recording sites, since any given stimulus was played to only a few recording sites (Schäfer et al., 2024). Models were implemented in PyTorch (version 2.9.0).

Several variants of the residual convolutional block, multi-site architecture led to similar neural prediction accuracy. The architecture and optimizer details of the model described in this paper are reported in Supplemental Table S1. The different hyperparameters tested include number of units (10-500), number of layers (2-12), nonlinearity in the final layer (double exponential, ReLU, leaky ReLU, and sigmoid), number of input frequency channels (32 and 72), number of gradient accumulation steps, and optimization parameters such as regularization (separately for shared dense layers, shared convolutional layers, and site-specific dense layers; range: 0-0.1).

Models were trained using the AdamW optimizer with dynamic gradient clipping (Loshchilov & Hutter, 2019). The loss function was normalized mean-squared error. Models were fit in three stages: 1) without the final-layer nonlinearity and with coarse training parameters (i.e., higher learning rate and gradient clipping threshold), 2) with the final-layer nonlinearity with coarse training parameters, and 3) with the final-layer nonlinearity with fine training parameters. At each stage, we used the first few epochs as warm-up epochs (He et al., 2016) and early stopping was used to prevent overfitting (Prechelt, 1998).

### Data analysis

#### Cross-validated PCA (cvPCA)

To estimate cvPCA, which offers unbiased estimates of dimensionality and variance explained between two datasets (e.g., recorded and predicted population PSTHs), we split the validation-stimulus trials (at least 10) into odd and even trials, and estimated averages for each split (Stringer et al., 2019). A PCA was fitted either to the even- or odd-trial data; for each fit, we projected both odd and even halves to the PCA and estimated the covariance between halves for each dimension. The average of both fits was used for the final covariance measure. cvPCA analysis between neural data and model predictions followed the same steps, except for each fit, we used model predictions instead of the other half of neural data. Prior to applying PCA, data were standardized (mean 0 and variance 1). The slope of the cvPCA eigenspectrum (i.e., variance explained [*v*] across PC dimensions [*n*]) was estimated as the exponent of a power-law function: *v* = *An^m^*, where *A* accounts for the DC offset in the eigenspectrum.

#### Representational Similarity Analysis (RSA)

To compare representational similarity across animals, we fit separate instances of the ACNet model for each animal. From these models, we obtained the manifold and predicted PSTHs for each animal. For RSA analysis, we first computed the representational dissimilarity matrix (RDM) for each data type (manifold, predicted PSTHs, and recorded PSTHs) for each animal. The RDM value at time points t1 (row index) and t2 (column index) was the Euclidean distance between the PCA-transformed activations for the two animals for each data type (Edelman, 1998; Kriegeskorte et al., 2008). PCA was used to keep 90% of the variance in the data for denoising. The representational similarity for two animals was quantified as the Pearson’s correlation coefficient (*r*) between their RDMs. We only used the upper-diagonal elements to estimate *r*, as the RDM is a symmetric matrix.

#### ESC-50 classification

The four classifiers applied to the ESC-50 data were trained using time-averaged embeddings of the following signals: recorded PSTHs, ACNet manifold activations, shuffled ACNet manifold activations (shuffled weights, intact temporal convolutional filters), and the gammatone spectrogram. For each classifier, we used the same architecture (a single hidden layer with 256 dimensions followed by an output layer). Classifiers were trained, validated, and tested with a five-fold jackknife. We conducted a neural architecture search over several hyperparameters (learning rate, weight decay, and dropout rate). We report the test results for the best validated classifiers.

#### Click-train simulations

Simulated PSTH responses to click trains by ACNet were used to classify neurons according to phase locking of their responses (Figure 5). Inter-click intervals (ICI) ranged from 3 ms to 100 ms. Each click was 100 µs in duration with a peak-to-peak amplitude of 1.58 (98 dB SPL). Stimuli were 1.5 seconds in duration and additionally included 250 ms of silence before and after the stimulus. The first 100 ms following stimulus onset were excluded from synchronization metrics (Lu et al., 2001).

To measure the strength of stimulus synchronization to click trains, we used vector strength (*VS*, Goldberg & Brown, 1969) and the Rayleigh statistic (RS, Mardia & Jupp, 2009). *VS* was defined as,

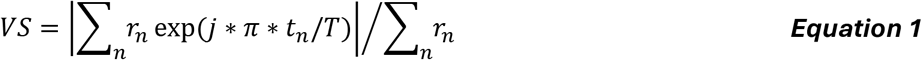

where *r_n_* and *t_n_* are the PSTH spike count and time (in seconds) at time bin n. T is the stimulus duration. *RS* is defined as 2*N* ∗ *VS*^2^, where N is the number of spikes within the time window. The ACNet PSTHs, which predict single-trial responses, were multiplied by a factor of 10 to obtain responses for 10 stimulus repetitions, as is typical of neurophysiological experiments (Lu et al., 2001). *VS* and *RS* were computed only for ICIs greater than or equal to 30 ms; for shorter ICIs, ACNet’s temporal resolution (10 ms) was inadequate to resolve individual clicks. The metric for rate coding, called discharge rate ratio or *DRR*, was defined as the ratio of the maximum discharge rate at ICIs less than 7 ms over maximum discharge rate at ICIs greater than 30 ms. Neurons were classified based on the following boundaries of these metrics: *RS*=13.8 and *DRR*=1.25. We used a more conservative (higher) criterion for *DRR* than previous studies (which used a DRR=1.0) as it better separated synchronized and nonsynchronized populations in our dataset, which may be due to a lack of stochastic noise in ACNet (Lu et al., 2001).

#### Manifold heterogeneity

Stimuli to probe heterogeneity of ACNet manifold activations (Figure 6) included pure tones (8 log-spaced steps from 250 Hz to 16 kHz), sinusoidal amplitude modulated (SAM) tones (two carrier frequencies: 1 and 8 kHz; two modulation frequencies: 5 Hz and 500 Hz), SAM octave-band noise (same carrier and modulation frequencies as SAM tones), upward and downward sweeps (250 Hz - 16 kHz for four durations: 25, 100, 400, and 1000 ms), and click trains (four inter-click intervals: 10, 20, 40, and 80 ms). Each stimulus was 500 ms in duration and presented at 55 dB SPL RMS sound level.

#### Spectrotemporal tuning subspace analysis

Tuning subspace analysis was used to characterize the spectrotemporal tuning properties of manifold dimensions (Figure 6C and 6D). This analysis reveals a set of spectrotemporal filters that fully describe the encoding subspace of each dimension (Wingert et al., 2026).

To measure the tuning subspace, we obtained activations to a subset of training stimuli for all ACNet manifold dimensions. At each time point, we estimated the dynamic spectrotemporal receptive field (dSTRF, Keshishian et al., 2020) as the gradient of the output (manifold activation) relative to the input gammatone spectrogram over the preceding 150 ms, a duration that covers the typical temporal integration window in AC (Sabat et al., 2025). We obtained a collection of these gradients at 4500 different time points spanning 10 min of natural stimuli, which is sufficient for subspace analysis to converge (Wingert et al., 2026). Then we applied PCA to estimate the most relevant spectrotemporal filters for each manifold dimension. For visualization of tuning across manifold dimensions, we only show the first principal component, which reflects its dominant tuning.

#### Frequency response area (FRA)

Tuning properties of manifold dimensions were estimated using FRA analysis, which reveals the response to tones of different frequencies and sound levels (Galambos & Davis, 1943; Sutter & Schreiner, 1991). The tone frequencies covered 200 Hz to 16 kHz in 24 log-scaled steps. Sound RMS level ranged from 20 dB SPL to 100 dB SPL in 5 dB steps. The stimulus duration was 1 second. For each stimulus, manifold responses were time averaged. Marginals of FRA were estimated as the RMS of activations across all frequencies (for level marginal) or sound levels (for frequency marginal) to quantify average tuning across sound level and frequency.

#### UMAP visualization for ESC-50 dataset

ACNet manifold activations were simulated for sounds in the ESC-50 dataset described above. For each stimulus (500 ms), manifold activations were time-averaged to obtain a 200-dimensional embedding vector. For visualizing the geometrical representation of these 50 categories in the manifold space, embeddings were projected to two dimensions using Uniform Manifold Approximation and Projection (UMAP) (McInnes et al., 2020). To identify the projection that best revealed category structure, we performed a grid search over two hyperparameters: the number of nearest neighbors (controls the balance between local and global structure; varied between 5 and 100) and the minimum inter-point distance in the low-dimensional space (controls cluster compactness; varied between 0 and .99). For each search, we selected 10 ESC-50 categories (two per taxonomic subgroup: Animals, Natural Soundscapes, Human non-speech, Interior, Exterior) that maximized a separation metric, which was the ratio of inter-category spread (mean pairwise Euclidean distance between category centroids) over within-category spread (mean distance of points to their centroid). Maximum separation was achieved for n_neighbors=50 and min_dist=0.5. These values were used in Figure 3 and Figure S1.

#### Temporal integration window analysis

We quantified how temporal integration window varies across layers of ACNet using the temporal context invariance analysis described in prior work (Norman-Haignere et al., 2022; Sabat et al., 2025). Natural sound segments of a fixed duration were concatenated to generate two different sequences with pseudorandom order so that each segment was preceded by two different segments (context). The similarity of responses of a unit to a segment in the two different contexts informs about the integration window (*T*) of that unit. A unit with *T* shorter than the segment duration produces increasingly similar responses over *T* following segment onset and identical responses after that. A unit with *T* longer than segment duration produces different responses, because its response still depends on the differing preceding context. One pseudorandom sequence pair was generated at each of six segment durations (15.625, 31.25, 62.5, 125, 250, and 500 ms); each sequence was 10 s in duration.

ACNet activations were obtained for all units across all six convolutional blocks. Since the model is deterministic, noise ceiling was 1 and no correction was applied to account for trial-to-trial variability that is present in neural data. Four of the six segment durations (15.625, 31.25, 62.5, and 125 ms) are not integer multiples of the model’s 10 ms output bin, so segment onsets fall at different phases within a response bin and lags cannot be referred to a common onset on the native grid. We therefore presented each sequence 16 times with the waveform advanced by 0.625 ms per repetition and interleaved the activations onto a 0.625 ms grid. For each unit and segment duration, we estimated CCC at each time lag relative to segment onset as the Pearson correlation between activations for the two sequence orders.

Following Sabat et al. (2025), we estimated the integration window by fitting a parametric window to the CCC. The window was a shifted, scaled Gamma density, 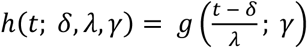 with 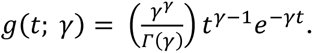. Because *δ* and *λ* are not themselves interpretable, the search was parameterized by two derived quantities: the window’s center (*c*), defined as the median of ℎ, and its width (*w*), defined as the smallest interval containing 75% of the mass of ℎ. Given a candidate window, the predicted CCC at each lag is the share of the response’s variance attributable to the shared segment, *P*(*τ*) = *s_s_*_ℎ*ared*_(*τ*)²/[*s_s_*_ℎ*ared*_(*τ*)² + *Σ_n_s_n_*(*τ*)²], where the sum runs over the surrounding segments. Parameters were fit per unit by a grid search over *γ* ∈ {1, …, 5}, 100 logarithmically spaced widths from 7.8 to 500 ms, and 100 logarithmically spaced centers from 1 to 250 ms, excluding combinations that violated causality of the window. The search minimized squared prediction error over lags from segment onset to segment offset at all six durations, with each duration weighted equally. The integration period was quantified as the fitted width.

#### Neuronal cell type classification

We assessed whether a unit’s physiological identity, defined by its joint spike-width × cortical-layer label (spike width: narrow vs. regular; layer: supragranular, granular, or infragranular, assigned from recording depth), could be decoded from the ACNet manifold. The analysis was restricted to A1/PEG units that had a response signal-to-noise ratio (SNR) > 0.1, similar to previous studies (Hamersky et al., 2025).

We used a two-layer perceptron (fully connected layers with LayerNorm, ReLU, and dropout) as the classifier, with manifold loading as the input and the physiological identity as output for each neuron. Classifiers were trained, validated, and tested with a site-based five-fold jackknife, with recording sites binned to balance unit counts across folds. We conducted a classifier architecture search over hidden-layer sizes, dropout rate, weight decay, and label-smoothing. For each feature set, the top validated configurations were each retrained with five random initializations, and the model with the highest mean cross-validated macro-F1 was retained. We report the test-set macro-F1 and accuracy of this best validated classifier.

### Statistics

For pairwise comparisons, we performed a Wilcoxon signed-rank test. For testing whether a metric monotonically increased or decreased across ACNet layers, we used Spearman rank-order correlation. Significance was determined at the α = 0.05 level after correcting for multiple comparisons using Bonferroni correction. These tests were performed in Python.

#### Bayesian models

We used Bayesian linear mixed-effects models (package brms::brm (Bürkner, 2017)) to assess how manifold-to-neuron loading similarity (MLS) for pairs of neurons varied across cortical regions (A1 and PEG), and whether MLS depended on their site identity (same-site, different sites in the same animal, and different sites in different animals) within each region (Equation 2). We also assessed whether MLS varied across cell types (narrow vs. regular spiking), and cortical layers (infra-granular, granular, and supra-granular), as described in Equation 3. All Bayesian statistical models were constructed in R (version 4.3.3).

#### Cortical area and recording site

The dependence of MLS on location within AC was modeled,

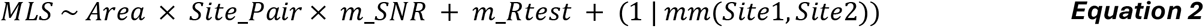

where *Site*_*Pair* distinguished same-site, cross-site same-animal, and cross-site different-animal pairs; *m*_*SNR* and *m*_*Rtest* are standardized geometric-mean SNR (as described above) and the standardized arithmetic-mean ACNet prediction accuracy for the two cells in each pair. *mm*(*Site*1, *Site*2) is a membership-model random intercept that accounts for site identity regardless of the order of a site pair. All same-area pairs (A1 and PEG) were included after SNR-based filtering and restricting pairs to within one octave of each other’s SNR. Because SNR was strongly correlated with prediction accuracy, we considered only pairs of neurons with similar SNR to simplify model design. Hypotheses about region (A1 vs. PEG) and connection type (same-site > cross-site) were evaluated as pairwise contrasts from posterior marginal means at mean SNR and Rtest (emmeans package; (Lenth, 2023)). We also tested a model that included pairs from different areas (A1 and PEG, Supplemental Table S3) to test whether MLS of pairs from across areas differed from MLS of pairs within an area. That model had the same form as Equation 2, with Area replaced by Area_pair (A1-A1, A1-PEG, or PEG-PEG).

#### Cortical layer and spike width

Models for assessing MLS across spike width (N, narrow; R, regular) and cortical layer (L1-3, supra-granular; L4, granular; L5-6, infra-granular) were constructed for either same-site or cross-site pairs:

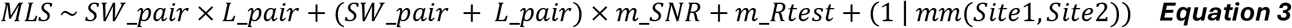

where *SW_pair* indicates combinations of spike widths (3 unique: RR, NN, RN) and *L_pair* indicates combinations of layers (6 unique). Spike-width-specific (NN > RR) and layer-specific (L13 > L4 > L56) hypotheses were evaluated as directional pairwise contrasts at *m_SNR* and *m_Rtest*. Cross-site pairs were subsampled to N = 150,000 as increasing N had little effect on statistical results. These results are detailed in Supplemental Tables S4 and S5, respectively.

Bayesian models used weakly informative priors (Normal(0, 0.5) for fixed effects, Normal(0, 0.2) for random-effect SDs), 4 chains × 6000 iterations (2000 warmup), and adapt_delta = 0.95. Model convergence was confirmed by ensuring the Gelman-Rubin diagnostic R-hat < 1.01 for all parameters.

#### Visualization of residuals for Bayesian models

To plot the similarity between pairs of cells (Figures 7 and S8), we removed the random effect of the recording sites and the fixed effect of Rtest (prediction accuracy) such that these plots only capture the residual effects to be consistent with our statistical models.

