## Supplemental material for "A low-dimensional, generalizable encoding manifold for auditory cortex"

#### Supplemental Figure S1

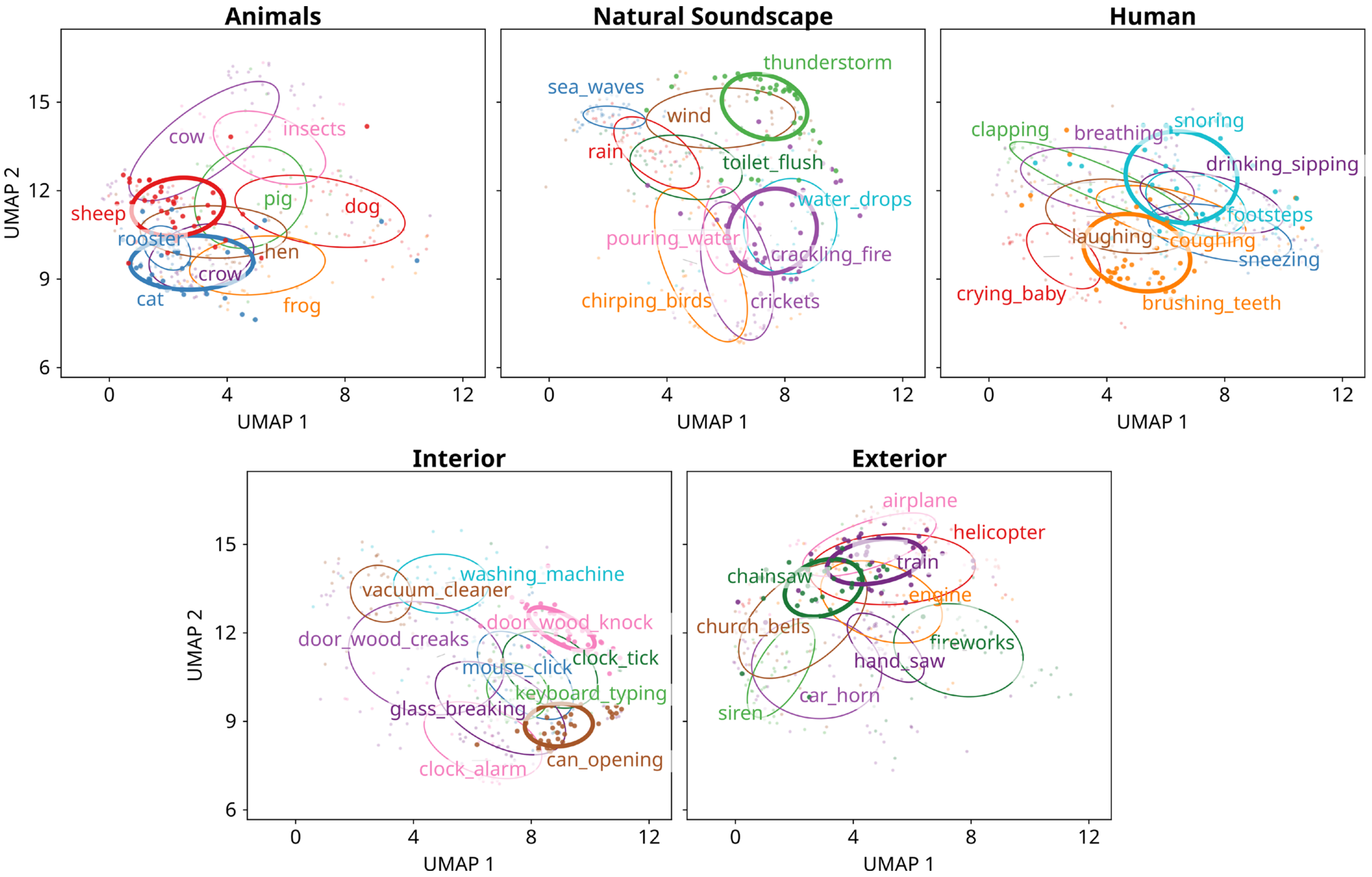

**Figure S1. UMAP representation in the ACNet manifold for all categories in the ESC-50 dataset.** Ellipses in individual panels highlight the 10 categories in each of the five super-groups of categories, all projected on the same axes. The 10 categories shown in Figure 3A are drawn with thick ellipses. Ellipses demarcate one standard deviation along the major and minor axes. The super-groups generally fall in different regions of the UMAP space (e.g., left for Animals, bottom right for Interior, upper left for Exterior, etc.).

#### Supplemental Figure S2

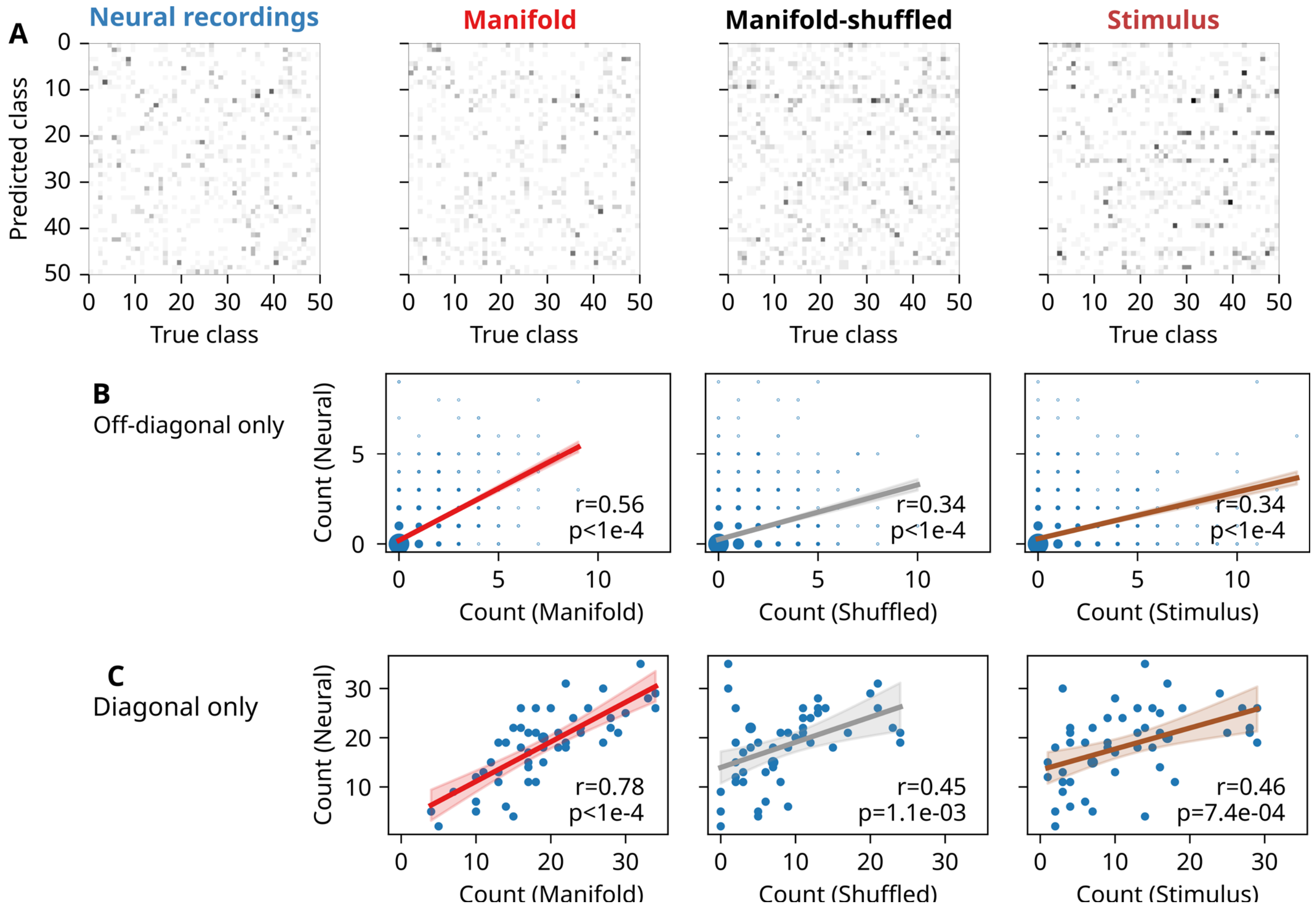

**Figure S2. Correlation of off-diagonal confusion matrices further highlights the neural alignment of the ACNet-based classifier.**

**(A)** Confusion matrices for the ESC-50 dataset, plotted as in Figure 3C, but with the diagonal elements masked to highlight the pattern of classification errors.

**(B)** Bubble plot showing correlation between off-diagonal elements, corresponding to incorrect predictions, in confusion matrices between neural and, in each panel, different non-neural models. Bubble size is proportional to the number of elements in that bin. Text denotes Pearson's correlation coefficient and p-value, and shading indicates 95% confidence interval for linear regression fit.

**(C)** Correlation of diagonal elements in confusion matrices from Figure 3C, plotted as in B.

#### Supplemental Figure S3

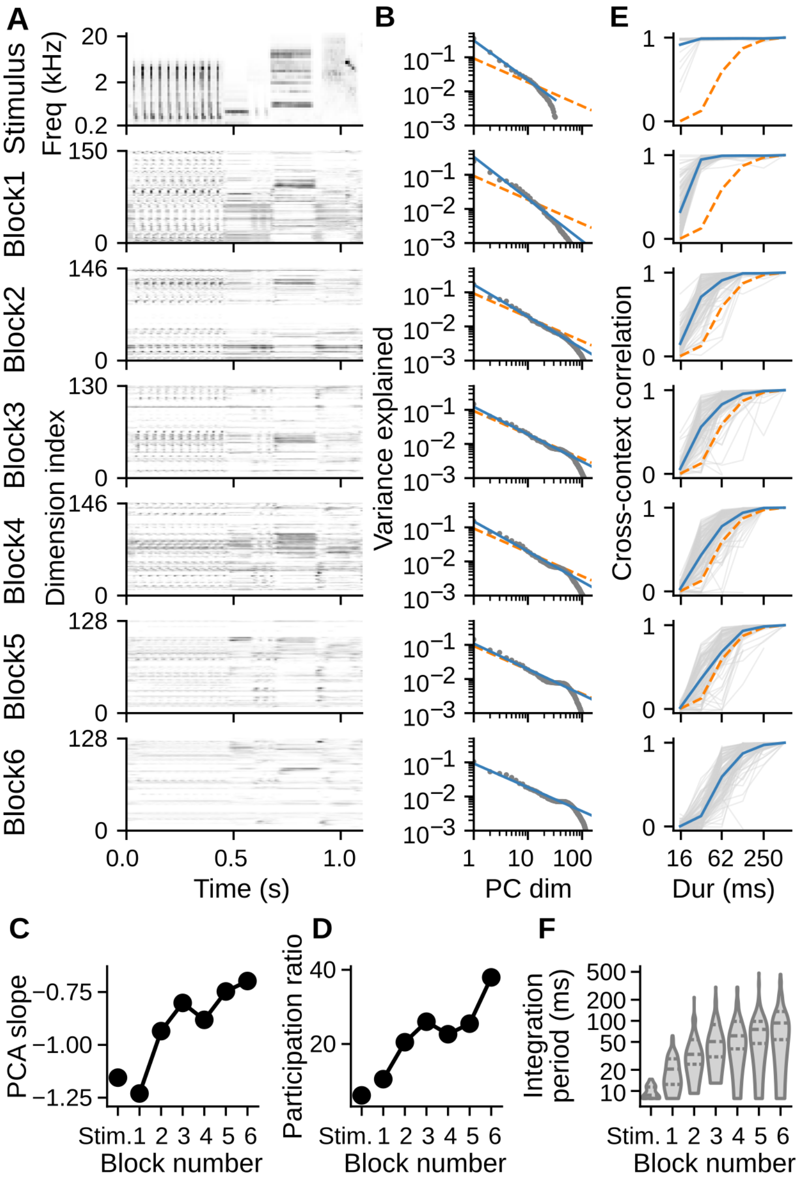

**Figure S3. Control model with flat architecture replicates key ACNet hierarchical properties, including increased efficiency and dimensionality, and longer, more diverse integration time constants.** Same format as Figure 4, but a model with 150 units in each of the blocks. This flat-architecture model performed slightly worse (median = 0.33) than ACNet (median = 0.34) in terms of predicting neural activity for the validation stimuli. Dead model units that were not activated by any stimulus were excluded from analysis. The model also showed higher slope (efficiency), higher participation ratio (dimensionality), and increased temporal integration (Spearman rank-order correlation: $r=0.65$, $p<1e-16$) and mean absolute deviation (MAD; Spearman rank-order correlation: $r=0.52$, $p<1e-16$) in CCC integration windows along the hierarchy. These results show that across-layer ACNet properties emerge from its hierarchy and do not depend on the differential number of units per layer.

#### Supplemental Figure S4

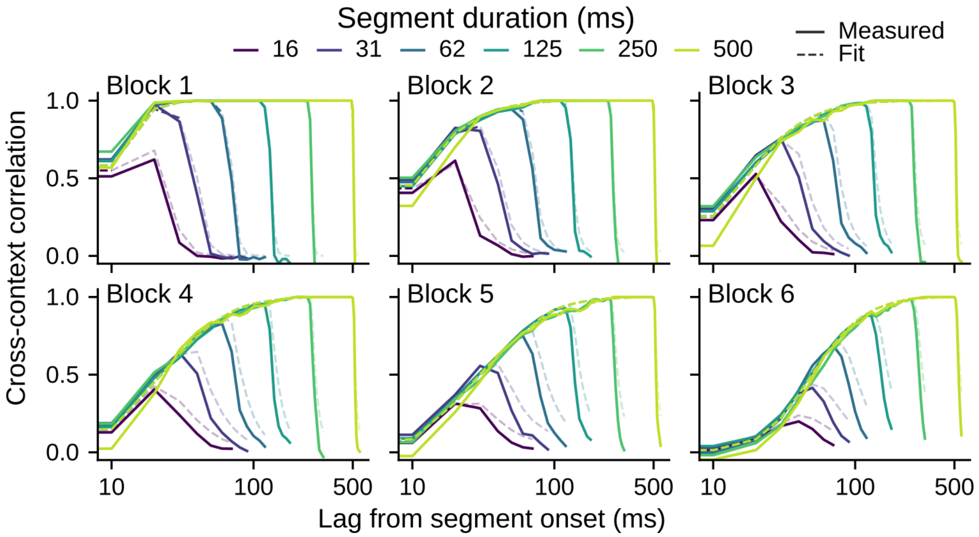

**Figure S4. Temporal context invariance analysis describes the temporal integration window for ACNet manifold units.** Solid lines show measured cross-context correlation (CCC) averaged across all units in each block for different stimulus durations (color). Dashed lines show the predicted CCC of the model that was used to estimate the temporal integration window.

#### Supplemental Figure S5

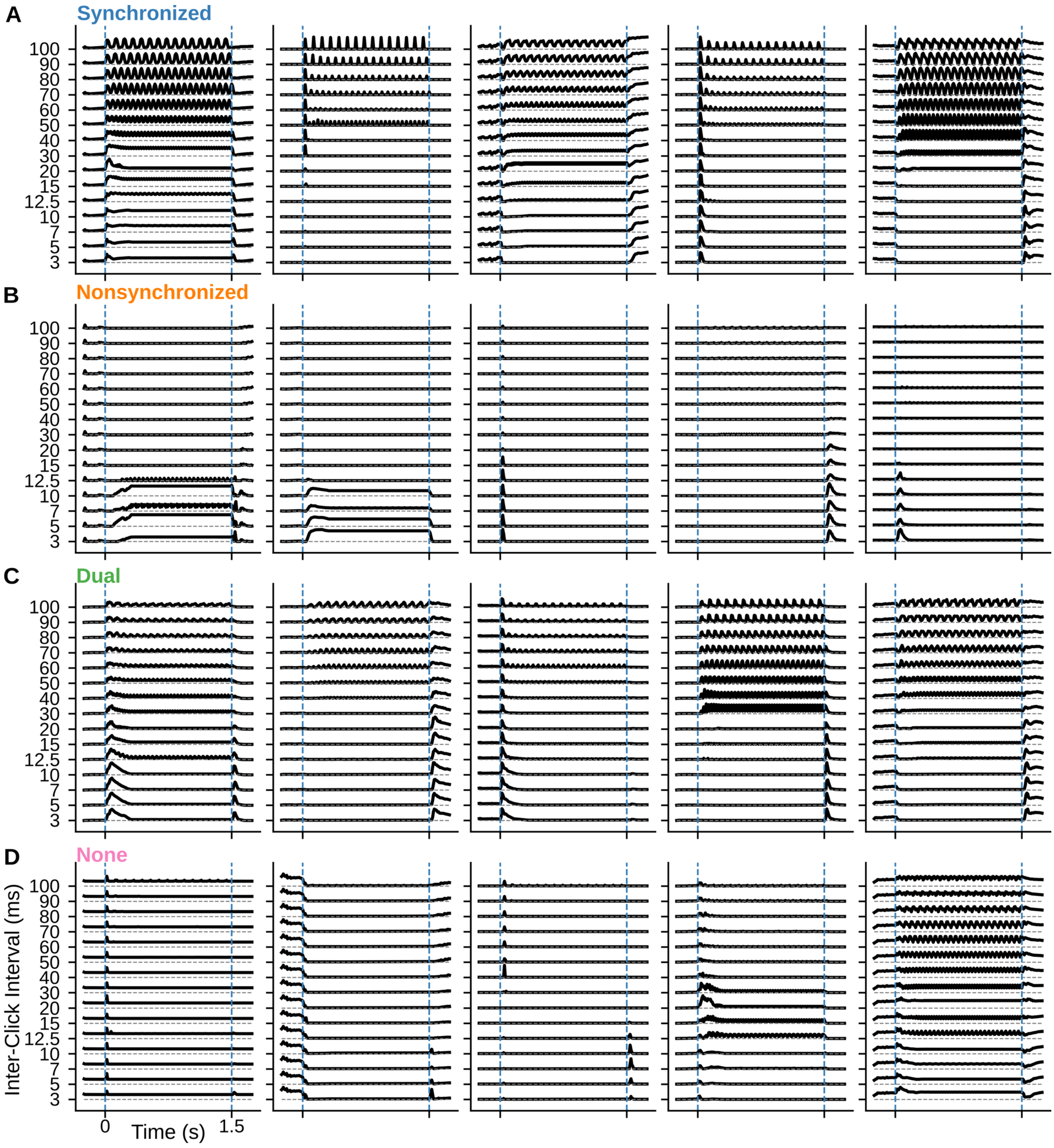

**Figure S5. Diversity of ACNet phase locking to click trains.** Each panel shows the simulated PSTH responses of a single unit to click trains of different inter-click intervals (ICI, 3 ms to 100 ms). Units were grouped according to their response pattern as in (Lu et al., 2001). Example neurons highlight the diversity of activity patterns in each group (5 units shown per group).

**(A)** For synchronized neurons, phase-locked responses could either be driven by excitation (column 2) or release from inhibition (column 5). Sustained responses may or may not be accompanied by an onset response.

**(B)** Responses of nonsynchronized neurons consisted of sustained firing without phase locking, onset or offset responses, and their combinations.
**(C)** Dual neurons showed properties of both synchronized and nonsynchronized neurons for fast and slow click trains, respectively.
**(D)** Neurons in the none category either displayed nonselective onset (column 1) or suppressive (column 2) responses, or band-limited tuning to ICI (columns 3-5).

#### Supplemental Figure S6

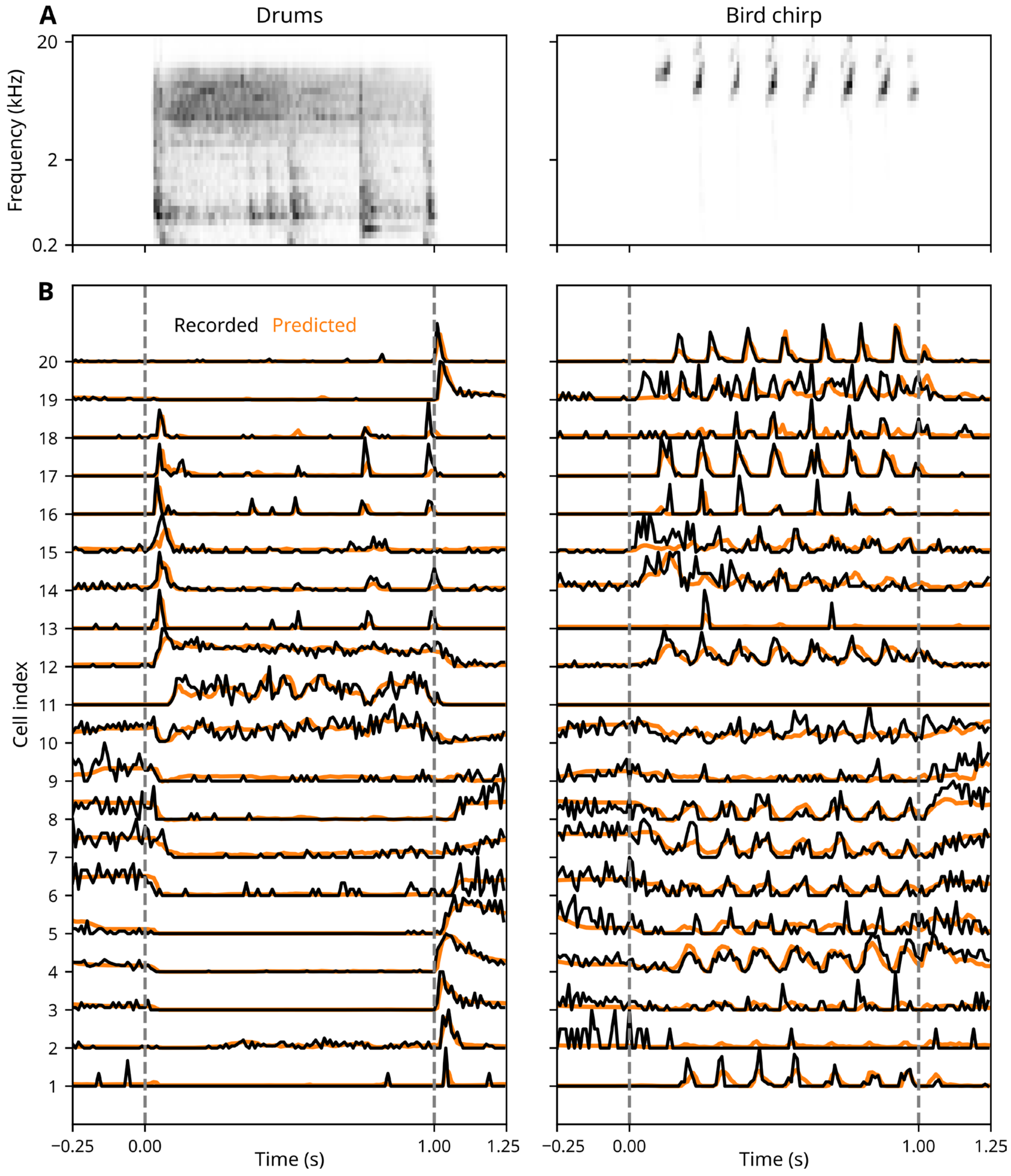

**Figure S6. ACNet captures diverse temporal patterns of response to natural sounds.**

**(A)** Gammatone spectrogram of two stimulus segments in the ACNet validation stimulus set (Drums, left, and Bird chirp, right).

**(B)** Recorded (black) and predicted (orange) PSTH responses to stimuli in A. Gray dashed lines indicate stimulus onset and offset. Neurons showed diverse response patterns consistent with the simulated click train responses, including non-synchronized onset, sustained, and offset responses for the drums segment, and synchronized responses to the chirp. Activity patterns also included sound-driven and sound-suppressed responses. ACNet predictions successfully reproduced this range of temporal response patterns.

#### Supplemental Figure S7

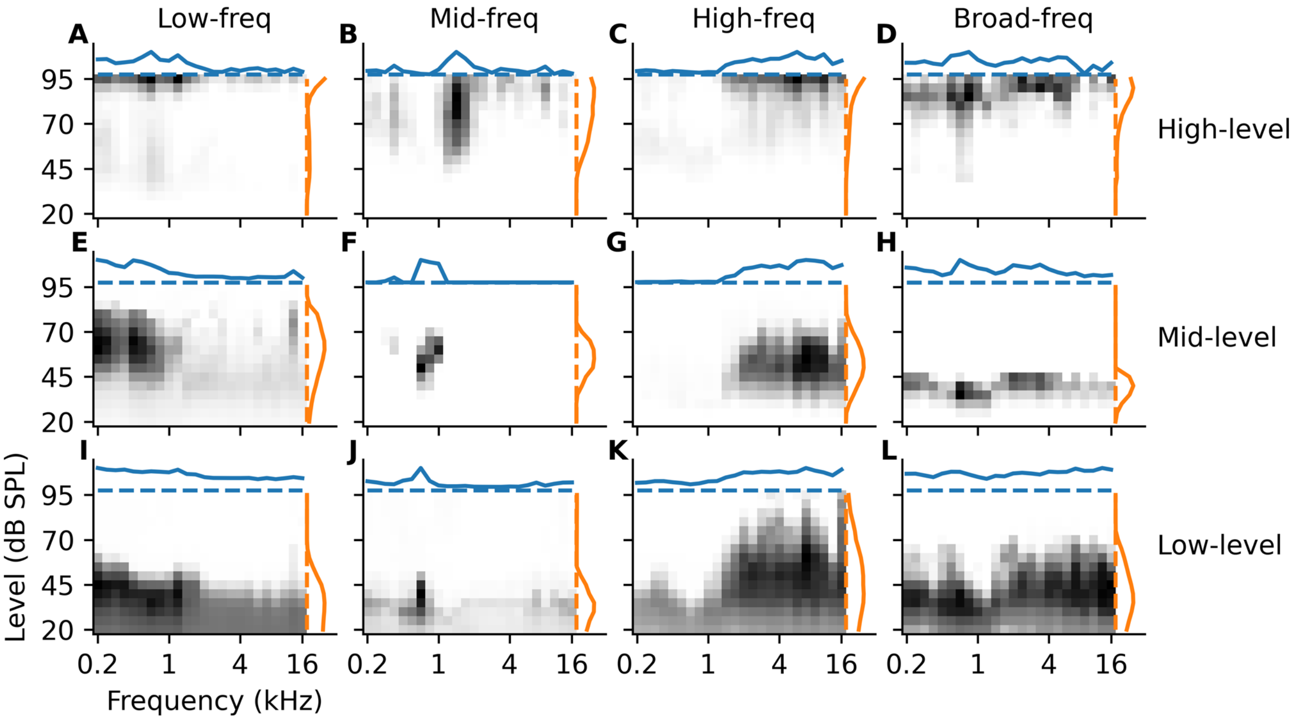

**Figure S7. FRA analysis shows a diversity of frequency tuning for manifold dimensions.** Each heatmap shows the time-averaged activation of a single manifold dimension per combination of tone frequency and level. Marginal frequency and level tuning, respectively, are plotted at the top (blue) and right (orange) of each panel. Dashed lines correspond to spontaneous activation. Frequency selectivity varies across columns: low-pass (A, E, I), band-pass (B, F, J), high-pass (C, G, K), and broad tuning (D, H, L). Sound level tuning varies across rows: high-pass (A-D), band-pass (E-H), and low-pass (I-L).

#### Supplemental Figure S8

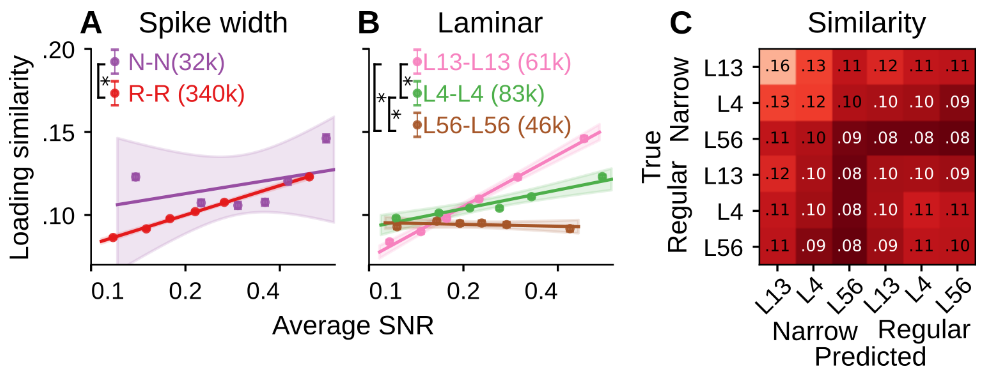

**Figure S8. Manifold loading similarity (MLS) followed similar but weaker trends for pairs of neurons across sites than within.** Comparison of response SNR versus MLS, plotted as in Figures 7D-7F, but for pairs of cells in different recording sites (pooled across same animal and different animals).

**(A)** MLS was greater for pairs of narrow-spiking (N) neurons than for pairs of regular-spiking (R) neurons ($\beta$ =-0.0089, 95% CI [-0.0124, -0.0053])**.**

**(B)** MLS decreased from supragranular (L13) to infragranular (L56) layers (L13 vs. L4: $\beta$ =0.0051, 95% CI [0.0024, 0.0077]; L4 vs. L56: $\beta$ =0.0083, 95% CI [0.0052, 0.0112]).

**(C)** Average MLS for pairs of neurons across cell and layer types. Detailed statistics are reported in Supplemental Table S5.

### Supplemental Table S1

**ACNet architecture and optimization hyperparameters.**

| Category | Parameter | Value / Detail |
|---|---|---|
| Input & Data | Input Frequency Channels | 32 |
| | Gammatone envelope compression | log10 (x) |
| | Temporal Resolution (fs) | 100 Hz |
| Architecture | Model Type | Multi-head Residual Convolutional Network |
| | Number of Blocks | 6 Blocks<br>- input block without skip connection<br>- Five residual convolutional blocks<br><br>Each block has<br>• Main branch with FIR filters + dense layer<br>• Skip connection (Dense) |
| | Hidden Dimensions | [75, 100, 125, 150, 175, 200] |
| | Temporal Kernel Duration | 70 ms (7 bins at 100 Hz) |
| | Final Layer Nonlinearity | Double exponential |
| Optimization | Optimizer | AdamW (with dynamic gradient clipping) |
| | Batch size | 25 stimuli |
| | Gradient Accumulation Steps | 8 |
| | Loss Function | Normalized mean-squared error |
| | L2 Weight Decay (Conv) | 0.0001 |
| | L2 Weight Decay (Shared Linear) | 0.0001 |
| | L2 Weight Decay (Site-specific) | 0.001 |

**Three-Stage Fitting Procedure**

| Parameter | Stage 1 (Linear) | Stage 2 (Coarse) | Stage 3 (Fine) |
|---|---|---|---|
| Nonlinearity | None | Double Exponential | Double Exponential |
| Initial Learning Rate | 0.001 | 0.001 | 0.0002 |
| LR Decay Factor (factor_lr) | 0.5 | 0.5 | 0.5 |
| Minimum Learning Rate (min_lr) | 0.001 | 0.0002 | 0.00001 |
| Loss Patience | 25 | 50 | 100 |
| Warmup Epochs | 20 | 20 | 25 |
| Early Stopping Patience | 50 | 200 | 1000 |

#### Supplemental Table S2

**Effect of site identity on MLS between pairs of neurons.** Site identity levels: same site, different sites in same animal, and different sites in different animals. Site_pair levels: SameSite; SameAnimal, different sites in the same animal; DiffAnimal, different sites in different animals (reference level, absorbed into the intercept). Coefficients and contrasts whose 95% interval excludes zero are shown in bold with an asterisk. m_SNR and m_Rtest are the pair's standardized geometric-mean SNR and arithmetic-mean ACNet prediction accuracy, as in Equation 2.

| Section / Parameter | Estimate | Error / Post. SD | 95% Interval (CI / HPD) |
|---|---|---|---|
| **1) Equation** | | | |
| $MLS \sim Area \times Site\_pair \times m\_SNR + m\_Rtest + (1 \mid mm(Site1, Site2))$ | | | |
| **2) Regression Coefficients** | | | |
| Parameter / Contrast | Estimate | Post. SD | 95% Interval (CI / HPD) |
| **Intercept** * | **0.0992** * | 0.0047 | [0.0900, 0.1086] |
| Area[PEG] | -0.0021 | 0.0110 | [-0.0239, 0.0197] |
| **Site_pair[SameAnimal]** * | **0.0074** * | 0.0008 | [0.0057, 0.0090] |
| **Site_pair[SameSite]** * | **0.0391** * | 0.0021 | [0.0351, 0.0432] |
| **m_SNR** * | **0.0060** * | 0.0006 | [0.0050, 0.0071] |
| **m_Rtest** * | **0.0189** * | 0.0005 | [0.0178, 0.0199] |
| Area[PEG] × Site_pair[SameAnimal] | -0.0072 | 0.0043 | [-0.0157, 0.0009] |
| Area[PEG] × Site_pair[SameSite] | -0.0015 | 0.0053 | [-0.0118, 0.0088] |
| Area[PEG] × m_SNR | 0.0029 | 0.0022 | [-0.0015, 0.0074] |
| **Site_pair[SameAnimal] × m_SNR** * | **0.0035** * | 0.0007 | [0.0020, 0.0049] |
| **Site_pair[SameSite] × m_SNR** * | **0.0108** * | 0.0019 | [0.0070, 0.0146] |
| Area[PEG] × Site_pair[SameAnimal] × m_SNR | 0.0038 | 0.0040 | [-0.0040, 0.0117] |
| Area[PEG] × Site_pair[SameSite] × m_SNR | 0.0029 | 0.0058 | [-0.0088, 0.0142] |
| **3) Contrasts – Site_pair per Area** | | | |
| Parameter / Contrast | Estimate | Post. SD | 95% Interval (CI / HPD) |
| A1 | | | |
| **(DiffAnimal) - (SameAnimal)** * | **-0.0074** * | - | [-0.0090, -0.0057] |
| **(DiffAnimal) - (SameSite)** * | **-0.0391** * | - | [-0.0431, -0.0351] |
| **(SameAnimal) - (SameSite)** * | **-0.0317** * | - | [-0.0359, -0.0276] |
| PEG | | | |
| (DiffAnimal) - (SameAnimal) | -0.0001 | - | [-0.0082, 0.0079] |
| **(DiffAnimal) - (SameSite)** * | **-0.0376** * | - | [-0.0472, -0.0279] |
| **(SameAnimal) - (SameSite)** * | **-0.0375** * | - | [-0.0490, -0.0262] |

#### Supplemental Table S3

**Effect of cortical area on MLS between pairs of neurons.** Data were either from primary (A1) or secondary (PEG) AC. Area_pair levels: A1-A1, PEG-PEG, and A1-PEG (reference level). Only different-site pairs are included. Coefficients and contrasts whose 95% interval excludes zero are shown in bold with an asterisk. m_SNR and m_Rtest are the pair's standardized geometric-mean SNR and arithmetic-mean ACNet prediction accuracy, as in Equation 2.

| Section / Parameter | Estimate | Error / Post. SD | 95% Interval (CI / HPD) |
|---|---|---|---|
| **1) Equation** | | | |
| $MLS \sim Area\_pair \times Site\_pair \times m\_SNR + m\_Rtest + (1 \mid mm(Site1, Site2))$ | | | |
| **2) Regression Coefficients** | | | |
| Parameter / Contrast | Estimate | Post. SD | 95% Interval (CI / HPD) |
| **Intercept *** | **0.0921 *** | 0.0049 | [0.0828, 0.1020] |
| Area_pair[A1-A1] | 0.0038 | 0.005 | [-0.0061, 0.0136] |
| Area_pair[PEG-PEG] | -0.0005 | 0.005 | [-0.0104, 0.0092] |
| Site_pair[SameAnimal] | 0.0016 | 0.0015 | [-0.0012, 0.0045] |
| **m_SNR *** | **0.0024 *** | 0.0011 | [0.0001, 0.0046] |
| **m_Rtest *** | **0.0200 *** | 0.0007 | [0.0187, 0.0213] |
| **Area_pair[A1-A1] × Site_pair[SameAnimal] *** | **0.0050 *** | 0.0022 | [0.0007, 0.0093] |
| Area_pair[PEG-PEG] × Site_pair[SameAnimal] | -0.001 | 0.0025 | [-0.0060, 0.0039] |
| Area_pair[A1-A1] × m_SNR | 0.0001 | 0.0014 | [-0.0026, 0.0029] |
| **Area_pair[PEG-PEG] × m_SNR *** | **0.0033 *** | 0.0017 | [3.8e-05, 0.0066] |
| Site_pair[SameAnimal] × m_SNR | 0.0009 | 0.0014 | [-0.0019, 0.0037] |
| Area_pair[A1-A1] × Site_pair[SameAnimal] × m_SNR | 0.003 | 0.0019 | [-0.0008, 0.0068] |
| **Area_pair[PEG-PEG] × Site_pair[SameAnimal] × m_SNR *** | **0.0074 *** | 0.0025 | [0.0026, 0.0123] |
| **3) Contrasts – Area_pair pairwise** | | | |
| Parameter / Contrast | Estimate | Post. SD | 95% Interval (CI / HPD) |
| (A1-PEG) - (A1-A1) | -0.006 | - | [-0.0156, 0.0036] |
| (A1-PEG) - (PEG-PEG) | 0.0009 | - | [-0.0088, 0.0105] |
| (A1-A1) - (PEG-PEG) | 0.0069 | - | [-0.0116, 0.0261] |

#### Supplemental Table S4

**Effect of cell type (regular [R] vs. narrow [N] spike width) and laminar identity (supragranular [L13], granular [L4], and infragranular [L56]) on MLS of pairs of neurons within the same recording site.** SW_pair levels: RR (reference), NR, NN. L_pair levels: L13-L13 (reference) through L56-L56. Coefficients and contrasts whose 95% interval excludes zero are shown in bold with an asterisk. m_SNR and m_Rtest are the pair's standardized geometric-mean SNR and arithmetic-mean ACNet prediction accuracy, as in Equation 3.

| Section / Parameter | Estimate | Error / Post. SD | 95% Interval (CI / HPD) |
|---|---|---|---|
| **1) Equation** | | | |
| $MLS \sim SW\_pair \times L\_pair + (SW\_pair + L\_pair) \times m\_SNR + m\_Rtest + (1 \mid Site1)$ | | | |
| **2) Regression Coefficients** | | | |
| Parameter / Contrast | Estimate | Post. SD | 95% Interval (CI / HPD) |
| **Intercept *** | **0.1768 *** | 0.0075 | [0.1621, 0.1915] |
| **SW_pair[NR] *** | **0.0210 *** | 0.0076 | [0.0059, 0.0355] |
| **SW_pair[NN] *** | **0.1490 *** | 0.0143 | [0.1211, 0.1772] |
| **L_pair[L13-L4] *** | **-0.0556 *** | 0.0055 | [-0.0667, -0.0451] |
| **L_pair[L13-L56] *** | **-0.0678 *** | 0.0067 | [-0.0808, -0.0547] |
| **L_pair[L4-L4] *** | **-0.0461 *** | 0.0061 | [-0.0580, -0.0342] |
| **L_pair[L4-L56] *** | **-0.0540 *** | 0.0063 | [-0.0663, -0.0418] |
| **L_pair[L56-L56] *** | **-0.0429 *** | 0.0075 | [-0.0577, -0.0282] |
| **m_SNR *** | **0.0518 *** | 0.0045 | [0.0429, 0.0607] |
| **m_Rtest *** | **0.0144 *** | 0.0022 | [0.0100, 0.0188] |
| SW_pair[NR] × L_pair[L13-L4] | -0.0094 | 0.0092 | [-0.0272, 0.0087] |
| **SW_pair[NN] × L_pair[L13-L4] *** | **-0.0559 *** | 0.0170 | [-0.0893, -0.0234] |
| **SW_pair[NR] × L_pair[L13-L56] *** | **-0.0214 *** | 0.0108 | [-0.0428, -0.0004] |
| **SW_pair[NN] × L_pair[L13-L56] *** | **-0.1311 *** | 0.0206 | [-0.1713, -0.0905] |
| SW_pair[NR] × L_pair[L4-L4] | -0.0121 | 0.0105 | [-0.0326, 0.0084] |
| SW_pair[NN] × L_pair[L4-L4] | -0.0199 | 0.0206 | [-0.0605, 0.0203] |
| **SW_pair[NR] × L_pair[L4-L56] *** | **-0.0480 *** | 0.0105 | [-0.0685, -0.0273] |
| **SW_pair[NN] × L_pair[L4-L56] *** | **-0.0984 *** | 0.0193 | [-0.1363, -0.0608] |
| **SW_pair[NR] × L_pair[L56-L56] *** | **-0.0423 *** | 0.0107 | [-0.0628, -0.0214] |
| **SW_pair[NN] × L_pair[L56-L56] *** | **-0.1323 *** | 0.0187 | [-0.1695, -0.0954] |
| **SW_pair[NR] × m_SNR *** | **-0.0071 *** | 0.0028 | [-0.0126, -0.0016] |
| SW_pair[NN] × m_SNR | -0.0071 | 0.0047 | [-0.0165, 0.0020] |

| Section / Parameter | Estimate | Error / Post. SD | 95% Interval (CI / HPD) |
|---|---|---|---|
| **L_pair[L13-L4] × m_SNR *** | **-0.0243 *** | 0.0045 | [-0.0331, -0.0153] |
| **L_pair[L13-L56] × m_SNR *** | **-0.0369 *** | 0.0056 | [-0.0477, -0.0259] |
| **L_pair[L4-L4] × m_SNR *** | **-0.0283 *** | 0.0049 | [-0.0380, -0.0188] |
| **L_pair[L4-L56] × m_SNR *** | **-0.0394 *** | 0.0052 | [-0.0495, -0.0291] |
| **L_pair[L56-L56] × m_SNR *** | **-0.0362 *** | 0.0061 | [-0.0482, -0.0242] |
| **3a) Contrasts – SW_pair (N vs R)** | | | |
| Parameter / Contrast | Estimate | Post. SD | 95% Interval (CI / HPD) |
| RR - NR | -6.4e-05 | - | [-0.0056, 0.0061] |
| **RR - NN *** | **-0.0796 *** | - | [-0.0921, -0.0671] |
| **NR - NN *** | **-0.0796 *** | - | [-0.0918, -0.0677] |
| **3b) Contrasts – L_pair (L13 / L4 / L56)** | | | |
| Parameter / Contrast | Estimate | Post. SD | 95% Interval (CI / HPD) |
| **(L13-L13) - (L13-L4) *** | **0.0621 *** | - | [0.0541, 0.0702] |
| **(L13-L13) - (L13-L56) *** | **0.0829 *** | - | [0.0727, 0.0924] |
| **(L13-L13) - (L4-L4) *** | **0.0510 *** | - | [0.0414, 0.0604] |
| **(L13-L13) - (L4-L56) *** | **0.0751 *** | - | [0.0652, 0.0846] |
| **(L13-L13) - (L56-L56) *** | **0.0644 *** | - | [0.0520, 0.0763] |
| **(L13-L4) - (L13-L56) *** | **0.0207 *** | - | [0.0118, 0.0290] |
| **(L13-L4) - (L4-L4) *** | **-0.0111 *** | - | [-0.0192, -0.0034] |
| **(L13-L4) - (L4-L56) *** | **0.0129 *** | - | [0.0051, 0.0212] |
| (L13-L4) - (L56-L56) | 0.0023 | - | [-0.0084, 0.0137] |
| **(L13-L56) - (L4-L4) *** | **-0.0318 *** | - | [-0.0416, -0.0223] |
| (L13-L56) - (L4-L56) | -0.0078 | - | [-0.0169, 0.0013] |
| **(L13-L56) - (L56-L56) *** | **-0.0185 *** | - | [-0.0297, -0.0071] |
| **(L4-L4) - (L4-L56) *** | **0.0240 *** | - | [0.0155, 0.0332] |
| **(L4-L4) - (L56-L56) *** | **0.0133 *** | - | [0.0016, 0.0248] |
| (L4-L56) - (L56-L56) | -0.0107 | - | [-0.0213, 0.0003] |

#### Supplemental Table S5

**Same as Table S4, but for pairs of neurons in different sites.** Since MLS was similar between pairs of sites in the same animal and across animals, data were pooled across these conditions. Symbols as in Table S4; the site random effect is mm(Site1, Site2) rather than Site1. Coefficients and contrasts whose 95% interval excludes zero are shown in bold with an asterisk. m_SNR and m_Rtest are the pair's standardized geometric-mean SNR and arithmetic-mean ACNet prediction accuracy, as in Equation 3.

| Section / Parameter | Estimate | Error / Post. SD | 95% Interval (CI / HPD) |
|---|---|---|---|
| **1) Equation** | | | |
| $MLS \sim SW\_pair \times L\_pair + (SW\_pair + L\_pair) \times m\_SNR + m\_Rtest + (1 \mid mm(Site1, Site2))$ | | | |
| **2) Regression Coefficients** | | | |
| Parameter / Contrast | Estimate | Post. SD | 95% Interval (CI / HPD) |
| **Intercept *** | **0.1059 *** | 0.0040 | [0.0981, 0.1139] |
| **SW_pair[NR] *** | **0.0084 *** | 0.0022 | [0.0042, 0.0127] |
| **SW_pair[NN] *** | **0.0362 *** | 0.0043 | [0.0279, 0.0446] |
| **L_pair[L13-L4] *** | **-0.0054 *** | 0.0016 | [-0.0085, -0.0024] |
| **L_pair[L13-L56] *** | **-0.0076 *** | 0.0017 | [-0.0109, -0.0042] |
| **L_pair[L4-L4] *** | **0.0044 *** | 0.0017 | [0.0011, 0.0078] |
| L_pair[L4-L56] | -4.9e-05 | 0.0017 | [-0.0033, 0.0033] |
| L_pair[L56-L56] | 0.0015 | 0.0022 | [-0.0027, 0.0058] |
| **m_SNR *** | **0.0233 *** | 0.0012 | [0.0209, 0.0256] |
| **m_Rtest *** | **0.0175 *** | 0.0005 | [0.0165, 0.0186] |
| SW_pair[NR] × L_pair[L13-L4] | -0.0049 | 0.0026 | [-0.0099, 9.7e-05] |
| **SW_pair[NN] × L_pair[L13-L4] *** | **-0.0129 *** | 0.0050 | [-0.0227, -0.0029] |
| **SW_pair[NR] × L_pair[L13-L56] *** | **-0.0143 *** | 0.0027 | [-0.0196, -0.0091] |
| **SW_pair[NN] × L_pair[L13-L56] *** | **-0.0321 *** | 0.0051 | [-0.0418, -0.0220] |
| **SW_pair[NR] × L_pair[L4-L4] *** | **-0.0234 *** | 0.0029 | [-0.0291, -0.0178] |
| **SW_pair[NN] × L_pair[L4-L4] *** | **-0.0274 *** | 0.0061 | [-0.0395, -0.0154] |
| **SW_pair[NR] × L_pair[L4-L56] *** | **-0.0278 *** | 0.0026 | [-0.0329, -0.0227] |
| **SW_pair[NN] × L_pair[L4-L56] *** | **-0.0421 *** | 0.0053 | [-0.0525, -0.0319] |
| **SW_pair[NR] × L_pair[L56-L56] *** | **-0.0344 *** | 0.0032 | [-0.0406, -0.0281] |
| **SW_pair[NN] × L_pair[L56-L56] *** | **-0.0566 *** | 0.0062 | [-0.0686, -0.0443] |
| **SW_pair[NR] × m_SNR *** | **-0.0128 *** | 0.0007 | [-0.0142, -0.0114] |
| **SW_pair[NN] × m_SNR *** | **-0.0092 *** | 0.0013 | [-0.0118, -0.0068] |

| Section / Parameter | Estimate | Error / Post. SD | 95% Interval (CI / HPD) |
|---|---|---|---|
| **L_pair[L13-L4] × m_SNR *** | **-0.0108 *** | 0.0012 | [-0.0132, -0.0085] |
| **L_pair[L13-L56] × m_SNR *** | **-0.0074 *** | 0.0013 | [-0.0100, -0.0049] |
| **L_pair[L4-L4] × m_SNR *** | **-0.0083 *** | 0.0013 | [-0.0109, -0.0057] |
| **L_pair[L4-L56] × m_SNR *** | **-0.0144 *** | 0.0013 | [-0.0170, -0.0119] |
| **L_pair[L56-L56] × m_SNR *** | **-0.0129 *** | 0.0018 | [-0.0164, -0.0093] |
| **3a) Contrasts – SW_pair (N vs R)** | | | |
| Parameter / Contrast | Estimate | Post. SD | 95% Interval (CI / HPD) |
| **RR - NR *** | **0.0079 *** | - | [0.0065, 0.0094] |
| **RR - NN *** | **-0.0089 *** | - | [-0.0124, -0.0053] |
| **NR - NN *** | **-0.0168 *** | - | [-0.0203, -0.0132] |
| **3b) Contrasts – L_pair (L13 / L4 / L56)** | | | |
| Parameter / Contrast | Estimate | Post. SD | 95% Interval (CI / HPD) |
| **(L13-L13) - (L13-L4) *** | **0.0078 *** | - | [0.0056, 0.0101] |
| **(L13-L13) - (L13-L56) *** | **0.0143 *** | - | [0.0117, 0.0167] |
| **(L13-L13) - (L4-L4) *** | **0.0051 *** | - | [0.0024, 0.0077] |
| **(L13-L13) - (L4-L56) *** | **0.0119 *** | - | [0.0093, 0.0143] |
| **(L13-L13) - (L56-L56) *** | **0.0134 *** | - | [0.0101, 0.0167] |
| **(L13-L4) - (L13-L56) *** | **0.0064 *** | - | [0.0045, 0.0084] |
| **(L13-L4) - (L4-L4) *** | **-0.0027 *** | - | [-0.0049, -0.0007] |
| **(L13-L4) - (L4-L56) *** | **0.0041 *** | - | [0.0022, 0.0060] |
| **(L13-L4) - (L56-L56) *** | **0.0056 *** | - | [0.0027, 0.0083] |
| **(L13-L56) - (L4-L4) *** | **-0.0091 *** | - | [-0.0114, -0.0067] |
| **(L13-L56) - (L4-L56) *** | **-0.0023 *** | - | [-0.0043, -0.0003] |
| (L13-L56) - (L56-L56) | -0.0009 | - | [-0.0037, 0.0018] |
| **(L4-L4) - (L4-L56) *** | **0.0068 *** | - | [0.0045, 0.0090] |
| **(L4-L4) - (L56-L56) *** | **0.0083 *** | - | [0.0052, 0.0112] |
| (L4-L56) - (L56-L56) | 0.0015 | - | [-0.0011, 0.0041] |
